# Preventing acute neurotoxicity of CNS therapeutic oligonucleotides with the addition of Ca^2+^ and Mg^2+^ in the formulation

**DOI:** 10.1101/2024.06.06.597639

**Authors:** Rachael Miller, Joseph Paquette, Alexandra Barker, Ellen Sapp, Nicholas McHugh, Brianna Bramato, Nozomi Yamada, Julia Alterman, Dimas Echeveria, Ken Yamada, Jonathan Watts, Christelle Anaclet, Marian DiFiglia, Anastasia Khvorova, Neil Aronin

## Abstract

Oligonucleotide therapeutics (ASOs and siRNAs) have been explored for modulation of gene expression in the central nervous system (CNS), with several drugs approved and many in clinical evaluation. Administration of highly concentrated oligonucleotides to the CNS can induce acute neurotoxicity. We demonstrate that delivery of concentrated oligonucleotides to the CSF in awake mice induces acute toxicity, observable within seconds of injection. Electroencephalography (EEG) and electromyography (EMG) in awake mice demonstrated seizures. Using ion chromatography, we show that siRNAs can tightly bind Ca^2+^ and Mg^2+^ up to molar equivalents of the phosphodiester (PO)/phosphorothioate (PS) bonds independently of the structure or phosphorothioate content. Optimization of the formulation by adding high concentrations (above biological levels) of divalent cations (Ca^2+^ alone, Mg^2+^ alone, or Ca^2+^ and Mg^2+^) prevents seizures with no impact on the distribution or efficacy of the oligonucleotide. The data here establishes the importance of adding Ca^2+^ and Mg^2+^ to the formulation for the safety of CNS administration of therapeutic oligonucleotides.

## Introduction

Oligonucleotide therapeutics is a class of drugs with significant potential for developing disease-modifying treatments. To date, there are eighteen approved oligonucleotide therapeutics^1^, including antisense oligonucleotides (ASOs), gapmers, splice-switching oligonucleotides (SSOs), aptamers, and small-interfering RNAs (siRNAs)^2^. Therapeutic oligonucleotides are used to mediate gene expression in the central nervous system (CNS), and several ASOs^34,5^ and SSOs are approved.^6,7^ While ASOs have been clinically evaluated for decades^8,9^, several chemical configurations of siRNAs, including C16 conjugates^10,11^ and di-valent^12,13^, have recently demonstrated wide distribution and long-term efficacy in rodents, sheep, and NHP brains.

The safety of ASOs in CNS differs significantly between blockmers, like Nusinersen^3^, and GapmeRs, like Tofersen^4^, and is highly affected by the sequence and modification pattern. SSOs have a broad safety profile in the CNS, whereas gapmers have shown variable tolerance *in vivo* with one compound (Tofersen)^4,5,14^ approved. In contrast, many other gapmers fail due to low efficacy and toxicity. Short-term neurotoxicity of ASOs upon CNS administration has been reported, and changes in calcium oscillation correlate to the degree of toxicity.^15^ The observed toxicity can be sequence-dependent with the involvement of G-rich regions linked to the phenomena.^16^ Combining *in silico* and experimental screening can predict toxicity and minimize the advancement of highly toxic compounds *in vivo*.^16^ All clinically qualified gapmer ASOs have a phosphodiester (PO)/phosphorothioate (PS) mixed backbone^6,17,18^ because reducing the PS backbone is essential for increased tolerability.^19^

Oligonucleotides are short nucleic acids in which the PO or PS groups form the sugar-phosphate backbone of the nucleotide chain. Each sugar-phosphate backbone carries a negative charge at physiological pH, resulting in highly negatively charged molecules. In solution, DNA and RNA molecules preferentially bind to divalent cations^20–22^, like Mg^2+^ and Ca^2+^, due to their strong electrostatic interactions with negatively charged groups with tightly bound divalent cations reported in many crystal structures and necessary for forming active sites.^23,24^ For therapeutic oligonucleotides, PBS or artificial CSF (aCSF, a buffer closely representing biological CSF composition) is conventionally used for administration. Interestingly, no systematic studies have been performed to optimize the formulation of oligonucleotides, a different class of molecules for RNA interference (RNAi) with distinct chemical features. Thus, optimizing oligonucleotide formulation for safe and efficient delivery to the CNS is necessary.^25–27^ We hypothesized that introducing oligonucleotides built on a negatively charged backbone into the CNS could cause neurotoxicity by binding divalent cations from the CSF or intracellular stores.

Huntington’s disease (HD) is a neurodegenerative disorder caused by a CAG trinucleotide-repeat expansion in the exon 1. HD has been extensively explored as a target in oligonucleotide therapeutics, with multiple validated and optimized siRNAs and ASOs available.^28–32^ The ASO targeting huntingtin (Tominersen) failed in a human phase III trial due to toxicity, represented by increased neurofilament light (NfL) and ventricular volume.^6,18^ Thus, the study of the toxicity of therapeutic oligonucleotides for HD is a reasonable approach to finding safe treatments for a devastating disease that has substantial unmet medical needs.

We demonstrate that di-siRNA target *Htt* induces acute abnormal behaviors upon administration of high doses in awake and isoflurane-anesthetized animals. We identified the observed abnormal behaviors as seizures using electroencephalography (EEG) and electromyography (EMG) in awake mice. We demonstrate that adding high Ca^2+^ and Mg^2+^ divalent cations to aCSF (∼10x of natural levels) prevents acute neurotoxicity and does not affect mRNA and protein silencing in wild-type and HD animal models. Thus, optimization of oligonucleotide formulation for CSF administration represents a promising approach for enhancing the safety of this new class of drugs.

## Results

### CNS injections of di-valent siRNA induce dose-dependent acute neurotoxicity, measurable by electrophysiology

In the previously published *in vivo* mouse studies, di-valent siRNA (di-siRNA) was delivered to the CNS using tribromoethanol (TBE, Avertin) as an anesthetic.^12^ This non-pharmaceutical grade injectable anesthetic deeply anesthetizes the mice for 60-90 minutes.^33^ Due to the extended period of anesthesia, acute abnormal behaviors that occurred following intracerebroventricular (ICV) di-siRNA administration were not recognizable. In these studies, animals wake up without severe adverse events, and robust mRNA and protein silencing is observed.^12,34,35^ Transitioning to the use of isoflurane flow as the anesthesia method for ICV injections revealed acute unfavorable behaviors, including seizures.

To understand the siRNA-induced acute neurotoxicity revealed when using isoflurane, we screened multiple doses of previously published di-siRNA^HTT12^ **(Figure 1A)** targeting the messenger RNA (mRNA) of huntingtin (*Htt*). Wild-type FVB mice were used for these studies to eliminate the potential of confounding variables in neurotoxicity readouts with disease models. The mice were anesthetized under isoflurane, injected ICV into both lateral ventricles, and monitored for up to an hour during recovery **(Figure 1B).** We developed an acute tolerability score to track specific observable abnormal behaviors following di-siRNA delivery into the CSF, which may not represent the typical tonic-clonic seizure (**Figure 1C).** Abnormal behaviors were scored based on a severity index (1 = least, 4 = most), similar to previous studies reported.^15,16^ The least severe behaviors, including shaking, hunched posture, stiff tail, and rapid eye movement, were assigned 1 point, full body muscle rigidity, ataxia, and barrel-rolling were assigned 2 points, and popcorning and foaming at the mouth were assigned 3 points towards severity. The most severe behavior that could occur within the one-hour monitoring period was death, which was classified as 4 points toward severity. A total score of 20 represents the worst seizure severity observed during recovery, and 0 represents no observed seizures or other adverse events. The average acute tolerability scores for all animals are shown in **Table S2**.

**Figure 1.**
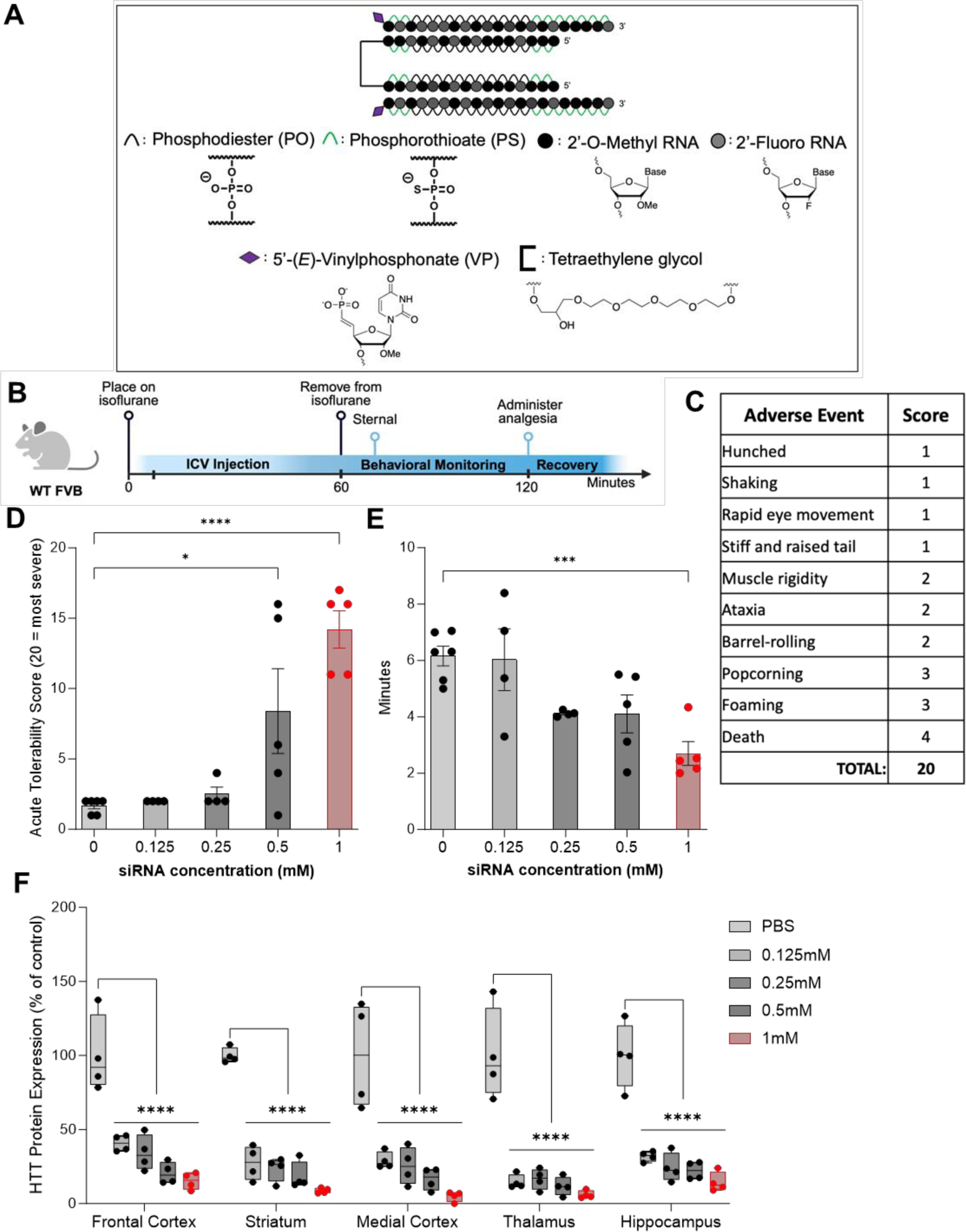
CNS injections of di-siRNA induce acute neurotoxicity in a dose-dependent manner. (A) The chemical structure of the di-siRNA^HTT^ used throughout this figure contains 74mM of PO/PS backbones, alternating 2’-OMe and 2’-Fluoro modifications, 5’-(E)-VP modifications, and a glycerol-tetraethyleneglycol linker. **(B)** Timeline showing the surgical procedure and recovery period for wild-type mice injected bilateral ICV with oligonucleotides under isoflurane anesthesia. (Created with BioRender.com) **(C)** Table outlining the acute tolerability scoring used to track adverse events that may not represent tonic-clonic seizures. The adverse events were scored based on an observational severity index (1 = least, 4 = most). A score of 20 represents the worst acute neurotoxicity observed during recovery, and a score of 0 represents no adverse events observed. **(D)** There was a dose-dependent increase in the severity of adverse events in wild-type mice injected with 0.125mM, 0.25mM, 0.5mM, or 1mM (∼28μg, ∼56μg, 112.5μg, and 225μg) di-siRNA^HTT^ in 10μL 1XPBS buffer. Administering the 0.5mM and 1mM di-siRNA^HTT^ resulted in significantly more severe abnormal behavioral phenotypes. Each data point represents one mouse (n= 4-6). *p=0.0163, ****p=<0.0001; data were analyzed using one-way ANOVA followed by Tukey’s post hoc test**. (E)** There was a significant difference in the time it took mice to be sternal following 1mM (225μg/10μL total dose) di-siRNA^HTT^ injections compared to the 1XPBS controls. Each data point represents one mouse (n= 4-6). ***p=0.0010.; data were analyzed using one-way ANOVA followed by Bonferroni’s multiple comparisons test**. (F)** Normal huntingtin (HTT) protein levels in di-siRNA^HTT^-treated mice versus 1XPBS controls measured by ProteinSimple Wes one-month post-ICV injections in the frontal cortex (FC), striatum (S), medial cortex (MC), thalamus (T), and hippocampus (H). There was significant HTT protein lowering across all brain regions measured with all four di-siRNA^HTT^ versus the 1XPBS vehicle control group (****p=<0.0001). Each data point represents one mouse and error bars represent mean values SEM (n=4). Data were analyzed using two-way ANOVA with Tukey’s multiple comparisons test.

ICV administration of di-siRNA^HTT^ at 0.125mM, 0.25mM, 0.5mM, and 1mM (∼28μg, ∼56μg, 112.5μg, and 225μg) resuspended in 10μL of 1X PBS revealed a dose-dependent neurotoxic readout. Mice injected with 0.125mM and 0.25mM di-siRNA^HTT^ had no further observable adverse events compared to the 1XPBS vehicle control mice **(Figure 1D)**. Mice injected with the vehicle control typically exhibited mild phenotypes, predominantly shaking and hunched posture. This was similar for mice injected with 0.125mM and 0.25mM di-siRNA^HTT^. Following administration of 0.5mM and 1mM di-siRNA^HTT^ injections, the severity of adverse events increased substantially, resulting in death for some animals at the maximum dose tested **(Figure 1D)**. Video recordings documented these findings. **(Supplemental Materials)** The mice injected with 0.5mM di-siRNA^HTT^ took significantly less time to right themselves and be sternal after being removed from isoflurane compared to the vehicle controls **(Figure 1E).** Higher doses of di-siRNA^HTT^ are necessary to reach >50% silencing of HTT protein.^12^ HTT protein levels were measured one-month following di-siRNA^HTT^ administration to confirm the dose-dependent effects **(Figure 1F).** The 1mM (225μg/10μL) di-siRNA^HTT^ delivered ICV provided maximum protein silencing (85-95% reduction across the brain), and based on this data, we chose to optimize the formulation of di-siRNA^HTT^ at the 1mM concentration and inject 225μg/10μL for these studies.

To test if the seizure-like behaviors described above are due to an interaction with the anesthetic or the oligonucleotide alone, *in vivo* Electroencephalography (EEG) and electromyography (EMG) were monitored during and after ICV di-siRNA^HTT^ injections in awake mice **(Figure 1A).** Wild-type mice were surgically implanted with EEG/EMG electrodes and a bilateral guide cannula targeting the lateral ventricles. ^36–40^ (Details in Methods) Following post-surgery recovery and an adaptation period to the recording conditions, awake mice were injected with di-siRNA^HTT^, and EEG/EMG was simultaneously recorded **(Figure 2A)**. Using SleepSign for Animal software (Kissei Comtec, Japan) assisted by spectral analysis using fast Fourier transform (FFT), polygraphic records were visually inspected offline for high amplitude EEG/EMG waves characteristic of seizures.^36–39^ Seizures were defined as periods with high frequency and high amplitude with a minimum duration of 5 seconds. Seizures were identified visually from EEG/EMG signals, and videos were checked to determine if there was a behavioral accompaniment. Mice injected ICV with 10μL 1X PBS as a control injection had no adverse events related to the injection and displayed typical EEG/EMG recordings throughout **(Figure 2C).** Representative EEG/EMG traces from a control mouse receiving an ICV injection of 1XPBS control showed no high-amplitude waves, indicating no seizures in response to the ICV injection **(Figure 2C).**

**Figure 2.**
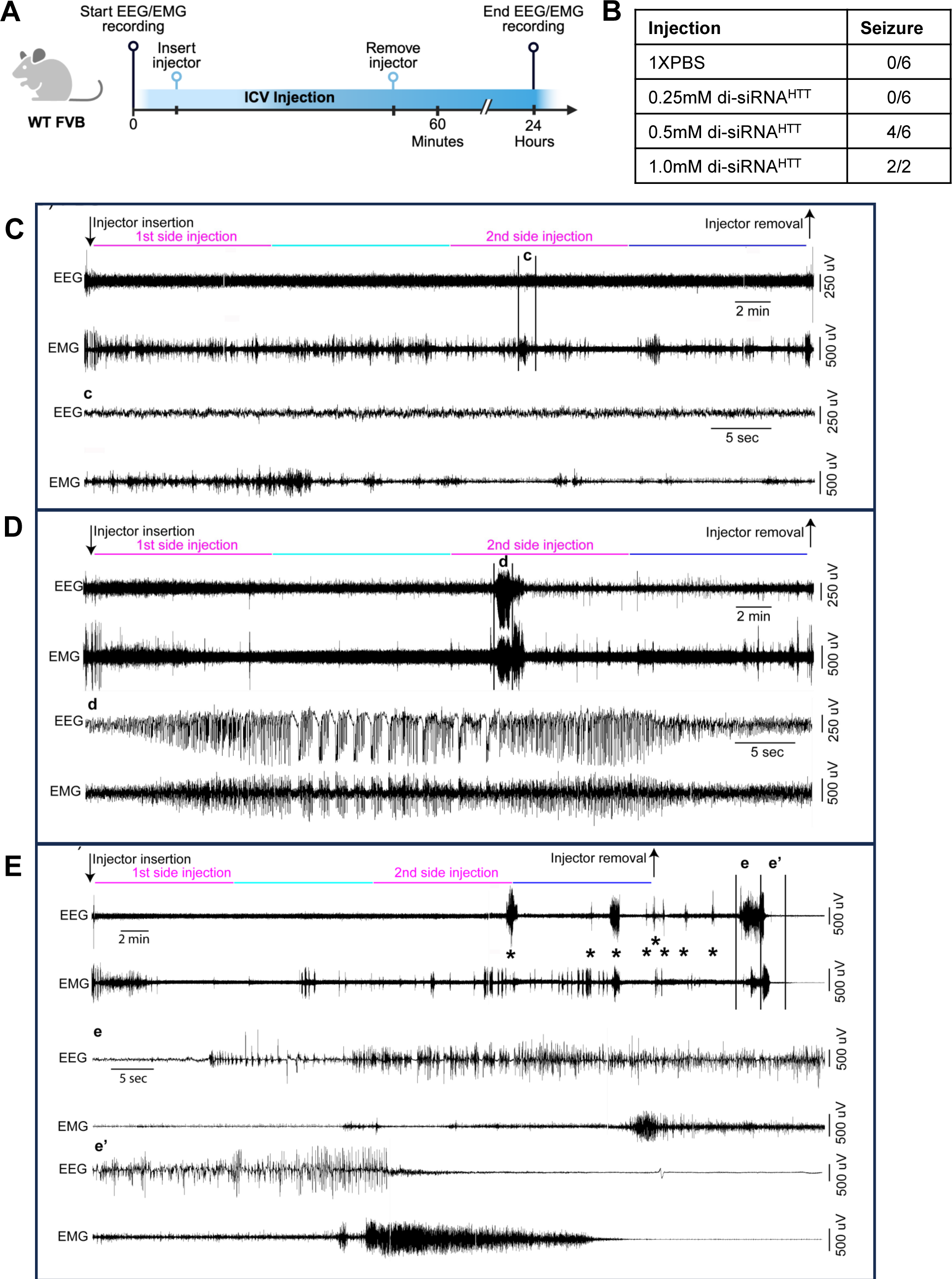
Electrophysiology defines the acute neurotoxicity induced by CNS injections of di-siRNA in awake mice. (A) Timeline showing the bilateral ICV injection of oligonucleotides associated with EEG/EMG recording in awake and freely moving wild-type mice. (Created with BioRender.com) **(B)** Table outlining the number of wild-type mice with seizure responses recorded by EEG/EMG following ICV injections of the following treatments. Treatments: 0.25mM (∼56μg) n=6, 0.5mM (112.5μg) n=6, or 1mM (225μg) n=2, di-siRNA^HTT^ in 10μL 1XPBS. **(C-E)** EEG/EMG examples from a freely moving mouse receiving bi-lateral ICV injection of 1XPBS (**C**), 0.5mM (112.5μg) di-siRNA^HTT^ **(D),** or 1mM (225μg) di-siRNA^HTT^ **(E)** in 10μL 1XPBS. The top left arrow indicates the insertion of the injector in the mouse guide cannula. The compounds were first injected in one of the two lateral ventricles and then in the second lateral ventricle (5μL in 10 mins, pink lines). Ten minutes separated the two side injections (light blue line). Following the end of the second side injection, the injector was left in place for an additional 10 minutes (dark blue line). The right arrow indicates the removal of the injector from the mouse guide cannula. (**c)** Zoom into the portion of **(C)** EEG-EMG example, delineated by the vertical bars. Note that the mouse injected with PBS displays normal EEG-EMG activity during the ICV injection. (**d**) Zoom into the portion of **(D)** EEG-EMG example, delineated by the vertical bars. The mouse injected with 0.5mM (112.5μg) di-siRNA^HTT^ 1XPBS displays a single seizure, characterized by high amplitude EEG waves and intense EMG activity, during the second side injection **(d)**. (**e**, **e’)** zoom into the portion of **(E)** EEG/EMG example, delineated by the vertical bars. The mouse injected with 1mM (225μg) di-siRNA^HTT^ **(E)** in 10μL 1XPBS displays multiple seizures, characterized by high amplitude EEG waves and intense EMG activity, starting around the beginning of the second side injection. Interestingly, about 10 minutes following the injector removal, the mouse enters a prolonged seizure **(e, e’)** that results in death, characterized by the absence of an EEG/EMG signal. Asterisks (*) denote seizures across the EEG/EMG example.

The acute tolerability of di-siRNA^HTT^ was evaluated with a dose-response of 0.25mM (∼56μg), 0.5mM (112.5μg), and 1mM (225μg) resuspended and injected in 10μL of 1X PBS into both lateral ventricles **(Figure 2B)**. Mice injected with 0.25mM (∼56μg/10μL) di-siRNA^HTT^ in 1XPBS had no observable effects and displayed EEG/EMG recordings similar to controls. Reaching higher di-siRNA^HTT^ concentrations of 0.5mM (112.5μg/10μL) elicited seizures in four out of six mice total, as recorded by EEG/EMG **(Figure 2D, d).** Representative EEG/EMG traces show striking differences between those from mice with normal phenotypes and those with tonic-clonic seizures **(Figure 2C-E)**. Two mice injected with 1mM (225μg/10μL) di-siRNA^HTT^ in 1XPBS died after displaying multiple seizures, characterized by high amplitude EEG waves and intense EMG activity, that started during the injection period. About 10 minutes following the removal of the injector, the mice entered a prolonged seizure **(e-e’)** that resulted in death, characterized by the absence of an EEG/EMG signal. Following a minimum of 24 hours of recording after the ICV injection, mice were deeply anesthetized [ketamine/xylazine, 200/20 mg/kg, intraperitoneal (IP)] and received bilateral ICV injections of blue dye (0.5μL per side). The brains were extracted and sectioned (antero-posterior axis, mid-line). Only data from brains showing dye evenly distributed throughout the CSF stores were included in the study **(Figure S2).** Video recordings were saved throughout the electrophysiology recording periods so overt behavioral phenotypes could be paired with adverse events recorded on EEG/EMG. These studies were invaluable for confirming the neurotoxic behavioral phenotypes observed following ICV injections in mice.

### Oligonucleotides can preferentially bind Ca^2+^ and Mg^2+^ divalent cations in an equimolar ratio to the PO/PS backbone *in vitro*

*In vitro*, we demonstrate that oligonucleotides can bind Ca^2+^ and Mg^2+^ divalent cations and retain these ions upon multiple water washes **(Figure 3)**. We used ion chromatography to gain insight into the binding capacity of cations to oligonucleotides **(Figure 3A).** In the sodium form following synthesis, di-siRNA^HTT^ with and without PS backbone modifications and a blunt version of di-siRNA^HTT^ **(Figure 3B)** were exposed to 100mM Ca^2+^, 100mM Mg^2+^, or 100mM Ca^2+^/Mg^2+^ solutions, then washed three times with sterile nuclease-free water and run on ion chromatography **(Figure 3B).** Representative chromatograph traces of the ion chromatography are displayed **(Figure 3C-E).** The standard traces are in gray, Mg^2+^ in blue **(Figure 3C),** Ca^2+^ in red **(Figure 3D),** and Ca^2+^/Mg^2+^ in green **(Figure 3E).** The separation of the cations is based on the size and charge of the ions interacting with the stationary phase of the column and the mobile phase. Changes in electrical conductivity are measured as peaks when the separated ions reach the detector, and the areas under the curves are relative to the specific cation concentrations compared to the standards. The results of the analysis are shown in **Figure 3F**.

**Figure 3.**
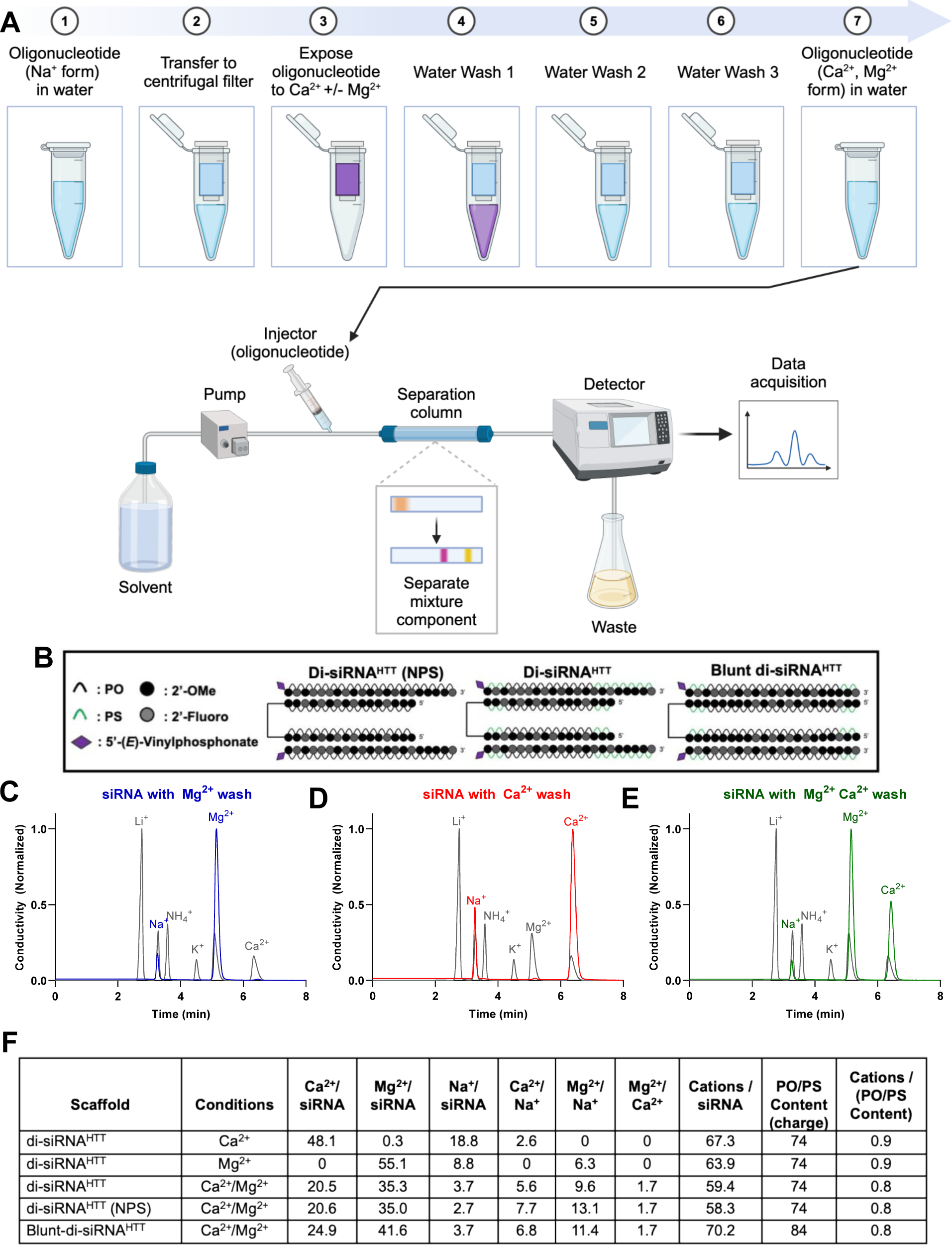
There is preferential binding of Ca^2+^ and Mg^2+^ divalent cations to the negatively charged PO/PS oligonucleotide backbone *in vitro*. (A) Schematic (Created with BioRender.com) depicting the ion chromatography workflow for *in vitro* divalent cation analysis of di-siRNAs. Oligonucleotides in Na^+^ form resuspended in water were transferred to centrifugal units, spun down to concentrate, and exposed to 100mM Ca^2+^ and/or Mg^2+^ solution. Following exposure to 100mM Ca^2+^ and/or Mg^2+^ the oligonucleotides were washed with water three times to remove any excess divalent cations in the solution. Oligonucleotides treated were then run through ion chromatography, and the cations of interest in the solution were quantified. The levels of each cation of interest were detected in real-time, and the data acquired was used to calculate the ratios in the table below. **(B)** The chemical structures of di-siRNAs analyzed using ion chromatography; previously described di-siRNA^HTT^, a di-siRNA^HTT^ with no PS modifications, and a blunt di-siRNA^HTT^ containing 10 more PS/PO backbones than di-siRNA^HTT^. Representative chromatograph traces for di-siRNA^HTT^ washed with 100mM Mg^2+^ **(C)**, 100mM Ca^2+^ **(D)**, or 100mM Ca^2+^ and Mg^2+^ **(E)**. **(F)** Table outlining the calculated ratios of oligonucleotide to cations derived from the ion chromatography experiments.

There was approximately a 1:1 ratio of the measured cations bound to phosphates (PO and PS) in the oligonucleotide backbone for every di-siRNA and cation exposure condition tested **(Figure 3F).** This may indicate high-affinity interactions, where one mole of the di-siRNA^HTT^ can bind up to 74 moles of cations to keep the charge neutrality of the solution **(Figure S3).** When di-siRNA^HTT^ was exposed to only Ca^2+^, the Ca^2+^ to Na^+^ ratio was approximately 3:1, suggesting that Ca^2+^ binds to phosphates 3X greater than monovalent Na^+^ cations **(Figure 3F)**. When di-siRNA^HTT^ was exposed to only Mg^2+^, the Mg^2+^ to Na^+^ ratio was approximately 6:1 **(Figure 3F)**. This suggests that Mg^2+^ has a 6X greater binding affinity to phosphates than Na^+^ and 2X greater binding affinity than Ca^2+^. At our saturation conditions, there was no detectable difference in binding capacity between PO and PS bonds **(Figure 3F).** A blunt-ended di-siRNA^HTT^ was also tested following exposure to 100mM Ca^2+^/Mg^2+^ to determine whether the molecule’s structure impacted the binding capacity of the divalent cations **(Figure 3D).** The blunt-di-siRNA^HTT^ contains ten more phosphates than the di-siRNA^HTT,^ which sequestered more cations in total but maintained the same approximate 1:1 ratio of backbone phosphates to cations. This data provides insight into the capacity of divalent cations to bind to oligonucleotides *in vitro*, which aids in the exploration of oligonucleotide-induced neurotoxicity and prevention.

### Adding Ca^2+^ and Mg^2+^ to aCSF mitigates acute neurotoxic behavioral phenotypes in a concentration-dependent manner

Here, we systematically evaluate if adding Ca^2+^ and Mg^2+^ to the formulation can eliminate acute neurotoxicity. Since Ca^2+^ phosphate salts have low solubility, mixing Ca^2+^ into the PBS buffer, specifically over time, results in the formation of micro-precipitates, which are severely toxic upon CSF administration.^19^ Due to this, a novel HEPES-based aCSF composition was used as a diluent. The final concentrations of the components are 137mM NaCl, 5mM KCl, 20mM D-+-glucose, and 8mM HEPES, with a final pH of 7.4. Natural CSF contains ∼ 2.2 mM Ca^2+^/Mg^2+^ levels in humans.^41,42^ The 1mM (225μg/10μL dose) di-siRNA^HTT^ (typical for ICV injection) **(Figure 4A)** has 74mM of negatively charged backbone. Theoretically, the oligonucleotide can bind 74 divalent cations equivalents, far exceeding ∼ 2.2 mM Ca^2+^/Mg^2+^ CSF levels in humans. Serial dilutions of Ca^2+^ and Mg^2+^ were made in the diluent aCSF (containing no Ca^2+^ and Mg^2+^) to make a range of divalent cation-enriched aCSF solutions. The stocks were later combined to create various ratios of Ca^2+^ and Mg^2+^ in aCSF.

**Figure 4.**
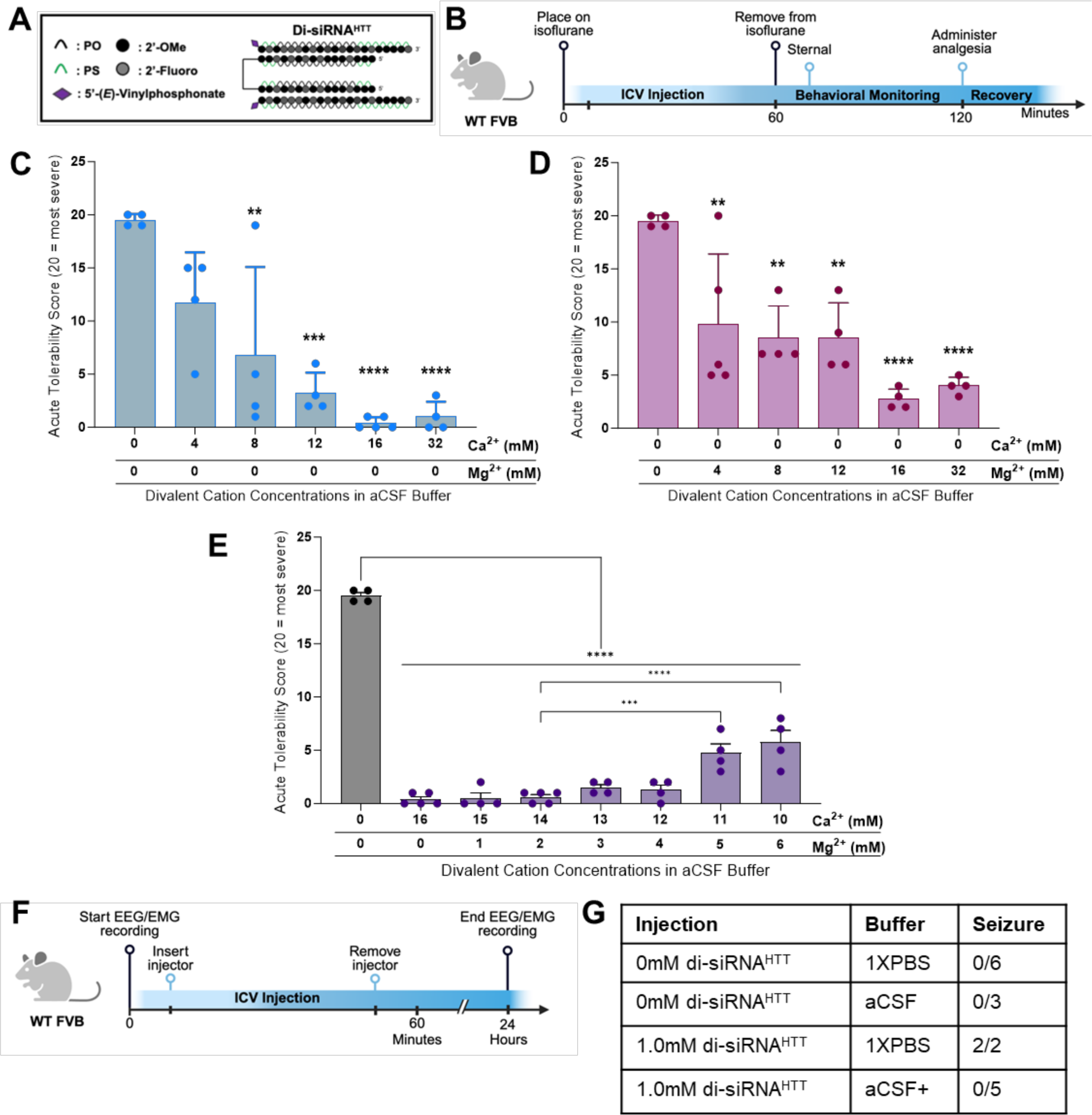
The acute neurotoxicity is preventable by adding Ca^2+^ and Mg^2+^ to aCSF buffer. (A) The chemical structure of di-siRNA^HTT^ used throughout this figure. **(B)** Timeline showing the surgical procedure and recovery period for wild-type mice injected bilateral ICV with oligonucleotides under isoflurane anesthesia. **(C)** Acute tolerability scores of wild-type mice injected with 1mM (225μg/10μL total) di-siRNA^HTT^ in aCSF with incrementally increased calcium concentrations. Increasing the Ca^2+^ concentration in aCSF buffer decreased the severity of adverse events in a dose-dependent manner. Each data point represents one mouse (n=4-6). **p=0.0018, ***p=0.0001, and ****p=<0.0001; data were analyzed using one-way ANOVA followed by Tukey’s post hoc test**. (D)** Acute tolerability scores of wild-type mice injected with 1mM (225μg/10μL total) di-siRNA^HTT^ in aCSF with incrementally increased Mg^2+^ concentration. Increasing the Mg^2+^ concentration in aCSF buffer significantly reduced the severity of adverse events in a dose-dependent manner. An excess of Mg^2+^ in aCSF increased hyperactivity in mice during recovery. Each data point represents one mouse (n=4-5). **p=0.0074 or **p=0.0038, and ****p=0<0.0001; data were analyzed using one-way ANOVA followed by Tukey’s post hoc test**. (E)** Acute tolerability scores of mice injected with 1mM (225μg/10μL total) di-siRNA^HTT^ in a range of aCSF buffers with incrementally increased Mg^2+^ concentration and reduced Ca^2+^ concentration. All seven aCSF formulations were significantly better tolerated than aCSF with no Ca^2+^ or Mg^2+^ (****p<0.0001). Decreasing the Ca^2+^ in aCSF to 11mM or 10mM with increased Mg^2+^ induced more severe adverse events (***p=0.0003 and ****p=<0.0001). Each data point represents one mouse (n=4-5).; data were analyzed using one-way ANOVA followed by Tukey’s post hoc test**. (F)** Timeline showing ICV injection procedure and EEG/EMG recording for wild-type mice injected bilateral ICV with oligonucleotides awake. (Created with BioRender.com) **(G)** Table outlining the number of wild-type mice with seizure responses recorded by EEG/EMG following intracerebroventricular injections of 0mM (vehicle control) or 1mM (225μg) di-siRNA^HTT^ in 10μL 1XPBS or aCSF+ (n=2-6 per treatment).

The acute neurotoxicity was evaluated with an escalation of Ca^2+^ concentration in aCSF buffer while maintaining the same 1mM (225μg/10μL total injected dose) di-siRNA^HTT^ in wild-type mice **(Figure 4B).** Increasing the Ca^2+^ concentration incrementally from 0mM to 16mM in aCSF buffer reduced the severity of seizures in a dose-dependent manner (n=4-6) **(Figure 4C).** The 1mM di-siRNA^HTT^ (225μg/10μL total injected dose) required 16mM Ca^2+^ in aCSF buffer to significantly reduce abnormal behaviors. Mice injected with excess Ca^2+^ (32mM in aCSF) did not have seizures but had prolonged recovery and sternal times **(Figure S4A).** We also tested a 16mM Ca^2+^ aCSF control with no siRNA to see if this amount of Ca^2+^ alone would be toxic. We found that injecting an excess of Ca^2+^ with no negatively charged siRNA resulted in prolonged sternal times but not seizures for some animals **(Figure S4A).** These data underlie the importance of having an exact ratio of Ca^2+^ to oligonucleotide in the injected compound to prevent neurotoxicity.

Due to the slight differences in binding capacities of Mg^2+^ and Ca^2+^ to the di-siRNA, we next evaluated the effect of adding Mg^2+^ into aCSF buffer while maintaining the same 1mM (225μg/10μL total injected dose) di-siRNA^HTT^. Similar to the Ca^+^ titration in aCSF buffer, increasing the Mg^2+^ concentration from 0mM to 16mM in aCSF buffer reduced the seizure phenotypes in a dose-dependent manner **(Figure 4D).** The same concentration of Mg^2+^ was required to improve the acute tolerability, compared to the di-siRNA^HTT^ delivered with Ca^2+^ in aCSF. Mice injected with excess Mg^2+^ (32mM in aCSF) had hyperactive phenotypes for approximately one hour following ICV injections. There was no difference in the time it took the mice to be sternal when injected with excess Mg^2+^ in aCSF compared to 1XPBS vehicle controls **(Figure S4B).**

To reach the necessary concentrations of divalent cations in aCSF and prevent delivering excess Ca^2+^ or Mg^2+^, we proposed that a combination of the two in aCSF may be the safest formulation. We screened 1mM (225μg/10μL total injected dose) di-siRNA^HTT^ formulated in aCSF buffer with increasing Mg^2+^ concentrations while reducing the Ca^2+^ concentration **(Figure 4E).** A range of aCSF buffers containing a concentration of 16mM total divalent cations were prepped, with differing Ca^2+^ and Mg^2+^ ratios. The buffer concentrations screened were 16:0, 15:1, 14:2, 13:3, 12:4, 11:5, and 10:6, Ca^2+^:Mg^2+^ respectively. Increasing the Mg^2+^ and decreasing the Ca^2+^ concentration was tolerated with 15:1 and 14:2 ratios of Ca^2+^:Mg^2+^ in aCSF **(Figure 4E).** Moving the Ca^2+^:Mg^2+^ ratio in aCSF to 13:3 and 12:4 was also tolerated, but mice experienced mild adverse events (shaking, hunched posture, and ataxia). Increasing the Mg^2+^ and decreasing the Ca^2+^ eventually produced observable toxicity **(Figure 4E).** These studies revealed that an ideal ratio of Ca^2+^ and Mg^2+^ divalent cations in aCSF to mitigate any adverse events completely was a 14:2 (14mM Ca^2+^ and 2mM Mg^2+^) ratio for 1mM di-siRNA^HTT^. Video recordings documented these findings. **(Supplemental Materials)** This 14:2 (14mM Ca^2+^ and 2mM Mg^2+^) formulation will be referred to as aCSF+.

To confirm the safety of formulating di-siRNA^HTT^ in aCSF+ before ICV injections, we studied EEG/EMG in wild-type mice. The same di-siRNA^HTT^ **(Figure 4A)** was injected bilaterally into both lateral ventricles of awake mice to confirm the absence of an acute seizure phenotype by EEG/EMG recording **(Figure 4F)**. As shown previously, ICV injections of 1mM (225μg/10μL total injected dose) di-siRNA^HTT^ in 1XPBS induced acute seizures **(Figure S5A).** Injecting 1mM (225μg/10μL total injected dose) di-siRNA^HTT^ in aCSF+ did not illicit any seizure activity recorded by EEG/EMG **(Figure S5B).** Video recordings were also populated throughout the electrophysiology experiments so overt behavioral phenotypes could be paired with adverse events recorded on EEG/EMG. Confirming the absence of spiking activity via electrophysiology is a precise measure of the safety of oligonucleotides delivered to the CNS. It should be considered as a safety readout for preclinical studies.

### Di-siRNA formulated in aCSF+ maintains efficacy and distribution throughout the mouse brain

To evaluate if the formulation of di-siRNA^HTT^ in aCSF+ may impact distribution and efficacy, *Htt* mRNA and HTT protein were measured two months following ICV injections. There was robust distribution and efficacy with 1mM (225μg/10μL total injected dose) di-siRNA^HTT^ formulated in aCSF+. The extent of *Htt* mRNA silencing is lower than HTT protein silencing because ∼50% of the mRNA is localized to the nucleus.^43^ As expected, di-siRNA^HTT^ formulated in aCSF+ reduced Htt mRNA by ∼50% **(Figure 5A)** and significantly reduced HTT protein expression by ∼90% across all brain regions evaluated **(Figure 5B, Figure S6)**. There was no change in GFAP mRNA levels between di-siRNA^HTT^ and vehicle controls **(Figure 5C).** The aCSF+ formulation developed to deliver oligonucleotides to the CNS safely does not impact the distribution in the brain or efficacy of mRNA and protein lowering.

**Figure 5.**
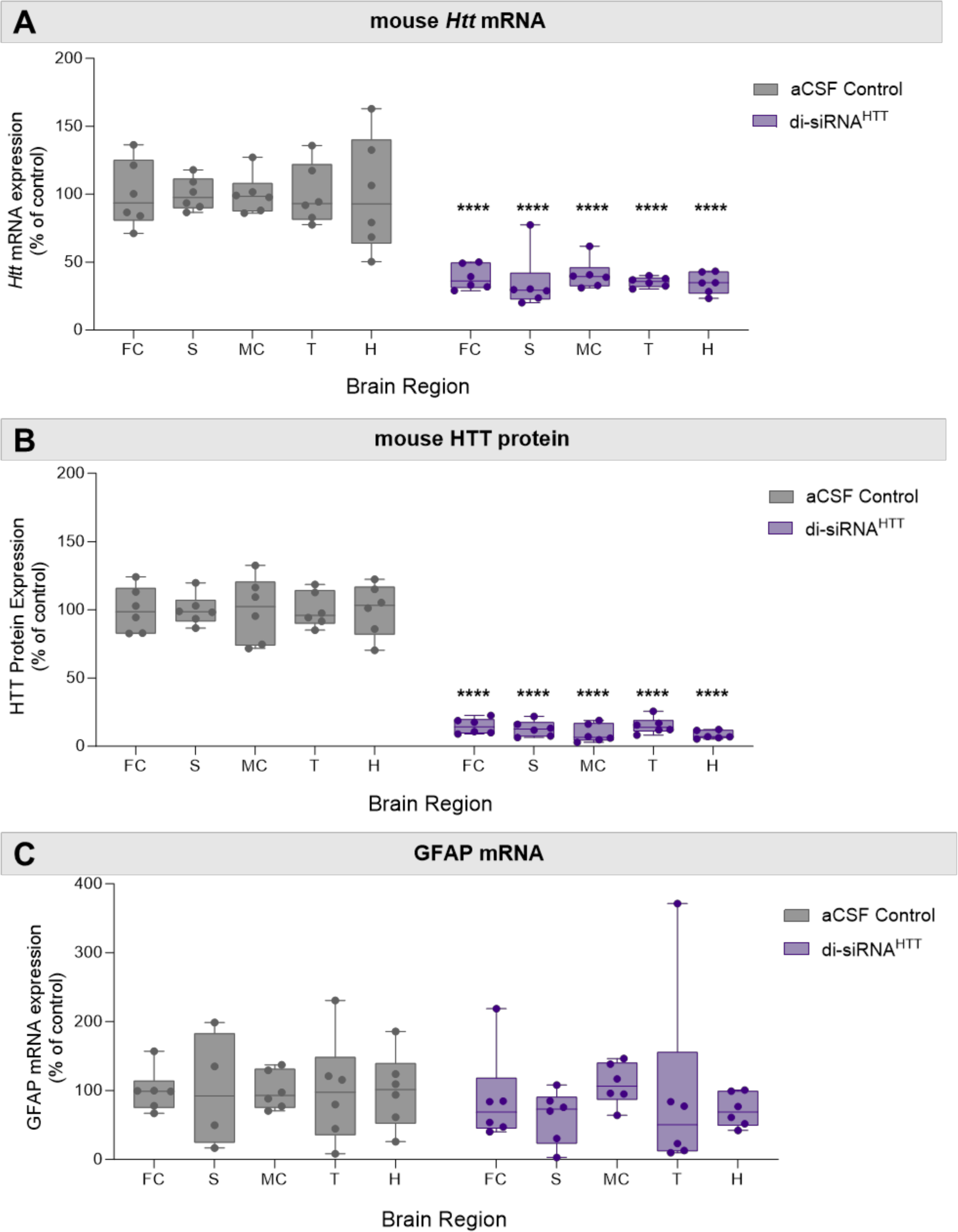
Di-siRNA^HTT^ formulated in aCSF+ maintains its distribution, efficacy, and safety in mouse brain. (A) Huntingtin *(Htt)* mRNA levels in the frontal cortex (FC), striatum (S), medial cortex (MC), thalamus (T), and hippocampus (H) were measured by Quantigen SinglePlex Assay. There was significant lowering in all brain regions two months following ICV injection of 1mM di-siRNA^HTT^ in aCSF+. **(B)** Mouse huntingtin protein (HTT) expression was measured by ProteinSimple Wes in the frontal cortex (FC), striatum (S), medial cortex (MC), thalamus (T), and hippocampus (H). Di-siRNA^HTT^ formulated in aCSF+ significantly reduced HTT expression in all brain regions evaluated two months following ICV injection compared to aCSF controls. **(C)** GFAP mRNA levels in the frontal cortex (FC), striatum (S), medial cortex (MC), thalamus (T), and hippocampus (H) were measured by Quantigen SinglePlex Assay. There was no change in GFAP mRNA levels in mice injected with 1mM di-siRNA^HTT^ in aCSF+ compared to aCSF controls. Each data point represents one mouse (n=6), and error bars represent mean values SEM. ****p<0.0001; data were analyzed using two-way ANOVA with Bonferroni’s multiple comparisons test.

### Maintaining the ratio of oligonucleotide to divalent cations in aCSF+ is safe at multiple doses

To test if the same aCSF+ buffer would mitigate acute neurotoxicity with higher doses of oligonucleotide, 450μg/10μL di-siRNA^HTT^ was injected. The same di-siRNA^HTT^ previously described **(Figure 6A)** was formulated in aCSF+ with a 2mM final concentration. Wild-type mice were injected bilaterally into both ventricles under isoflurane anesthesia and monitored post-operatively **(Figure 6B).** With a 450μg (2mM) di-siRNA^HTT^ dose, aCSF+ did not improve the acute tolerability. Doubling the Ca^2+^ and Mg^2+^ concentrations in aCSF+ to maintain a consistent ratio of di-siRNA^HTT^ and divalent cations was necessary to mitigate acute neurotoxicity. Interestingly, the 2mM (450μg/10μL dose) di-siRNA^HTT^ in aCSF+ containing only Ca^2+^ was not tolerated, whereas aCSF+ enriched with Ca^2+^ and Mg^2+^ completely mitigated acute neurotoxicity **(Figure 6C).** This indicated that both Ca^2+^ and Mg^2+^ may be required in aCSF+ in a consistent ratio to the oligonucleotide concentration in the solution. Safely injecting the maximum deliverable di-siRNA^HTT^ dose in mice is an important step forward for CNS oligonucleotide therapeutic development.

**Figure 6.**
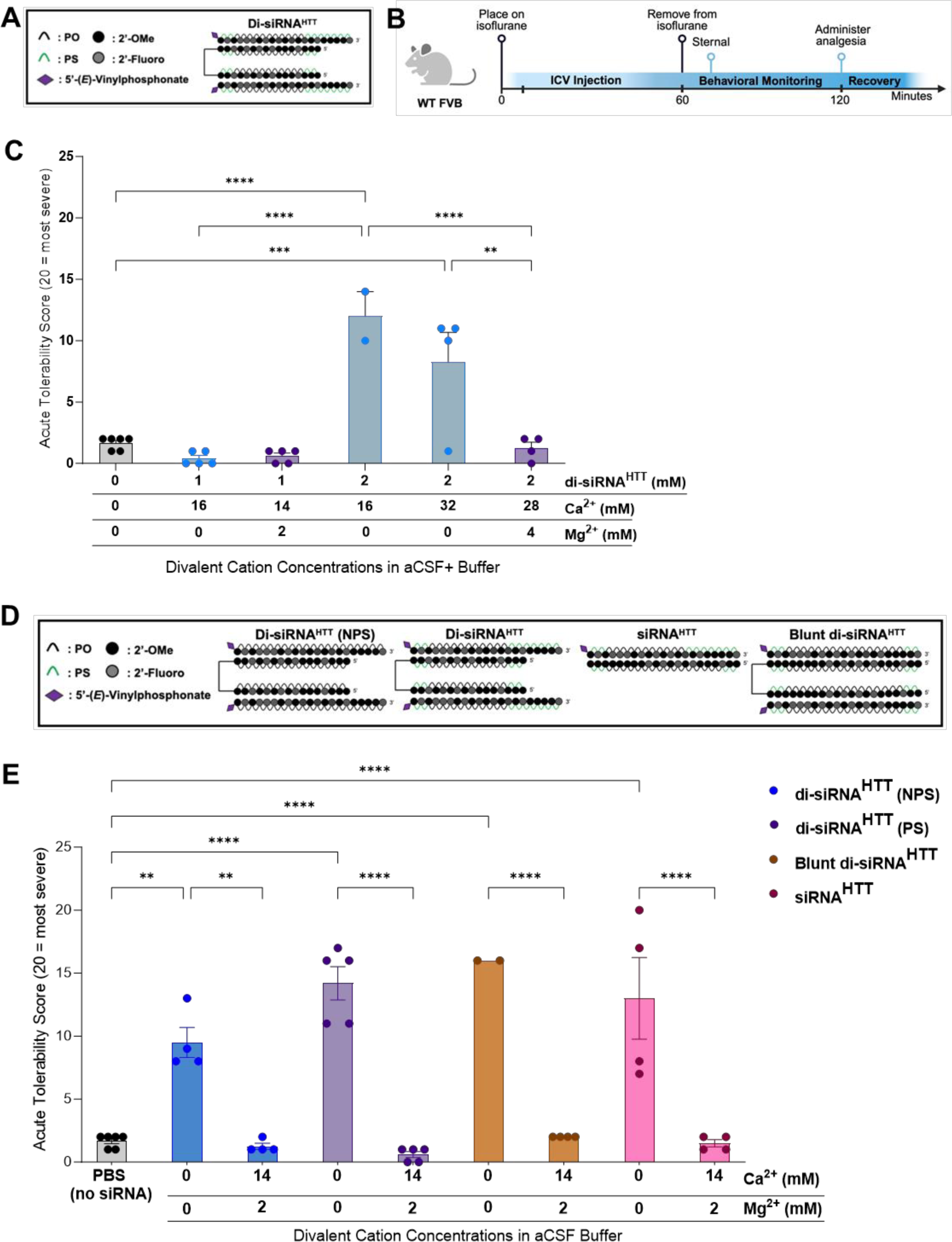
The new aCSF+ formulation is safe at multiple doses when the ratio of di-siRNA to divalent cations is consistent, with structure and backbone modifications having minimal impact on neurotoxicity. (A) The chemical structure of di-siRNA^HTT^ used throughout this figure. **(B)** Timeline showing the surgical procedure and recovery period for wild-type mice injected bilateral ICV with oligonucleotides under isoflurane anesthesia. (Created with BioRender.com) **(C)** Acute tolerability scores following ICV injection of 1mM (225μg/10μL total) and 2mM (450μg/10μL total) di-siRNA^HTT^ in aCSF+ with different concentrations of Ca^2+^ and Mg^2+^. There was no difference in tolerability in mice injected with 1mM di-siRNA^HTT^ in aCSF with either 16mM Ca^2+^ or 14mM Ca^2+^ and 2mM Mg^2+^. Injecting mice with 2mM di-siRNA^HTT^ in aCSF with 16mM Ca^2+^ was not well tolerated compared 1mM di-siRNA^HTT^ or aCSF controls (****p=<0.0001). Injecting 2mM di-siRNA^HTT^ in aCSF with 32mM Ca^2+^ was better tolerated, but still significantly worse than vehicle controls (***p=0.0002). Maintaining the same ratio of di-siRNA^HTT^ and divalent cations in aCSF+ significantly improved the acute tolerability (****p=<0.0001) and there was no difference compared to aCSF controls. **(D)** The chemical structures of siRNA oligonucleotides used in this figure: the previously described di-siRNA^HTT^ with PS modifications, di-siRNA^HTT^ (NPS) with 0 PS modifications, a monovalent siRNA^HTT^, and a blunt di-siRNA^HTT^. **(E)** Acute tolerability scores following ICV administration of the four different siRNAs in 1XPBS or aCSF+. The di-siRNA^HTT^ (NPS) was slightly better tolerated (**p=0.0011) than the di-siRNA^HTT^, siRNA^HTT^, and blunt di-siRNA^HTT^ (****p=<0.0001), with 1mM (225μg/10μL total) compared to the 1XPBS controls. Formulating the different siRNAs in aCSF+ significantly improved the acute tolerability (**p=0.0019 and ****p=<0.0001). There was no significant difference in the tolerability between the four di-siRNA^HTT^ scaffolds injected in aCSF+ compared to the 1XPBS controls. Each data point represents one mouse (n=2-6) with SEM. Data were analyzed using one-way ANOVA followed by Tukey’s post hoc test.

### Di-siRNA phosphorothioate content and structure have minimal impact on the divalent cation formulation necessary to prevent neurotoxicity

Throughout these studies, other variables within the oligonucleotide makeup that may contribute to acute neurotoxicity were considered. The addition of PS modifications to oligonucleotides is commonly used to enhance the stability and duration of effect.^44–46^ These are critical factors for their usefulness as therapeutics but come with the disadvantage of increased toxicities.^47–49^ To further understand the mechanism of binding between di-siRNA^HTT^ and divalent cations in CSF, wild-type mice were injected under isoflurane **(Figure 6B)** with 1mM (225μg/10μL dose) di-siRNA^HTT^ containing no PS content, and the previously described di-siRNA^HTT^ containing 26mM PS in 1XPBS or aCSF+ **(Figure 6D).** The severity of seizures observed was reduced with decreased PS content in di-siRNA^HTT^. The di-siRNA^HTT^ with 26mM PS content induced severe acute adverse events compared to 1XPBS vehicle controls **(Figure 6E).** When the same di-siRNAs were formulated in aCSF+ for ICV injection, nearly complete mitigation of all adverse events was observed **(Figure 6E).** The significant difference in acute tolerability represents that the oligonucleotide-induced neurotoxicity is not dependent on PS modifications, as seen in the cases of ASOs and hepatotoxicity.^47,48,50^

Since di-siRNAs are only one class of siRNA oligonucleotides in development for treating CNS diseases, we next assessed whether aCSF+ would be safe for other variations of HTT-targeting siRNAs. The di-siRNA^HTT12^ used for most of the studies presented here comprises a dimeric sense strand, or two 16-mer sense strands linked together, each bound to a 21-mer antisense strand targeting *Htt* mRNA. The longer antisense strand creates an overhang region containing single-stranded nucleic acids, which may influence this neurotoxic seizure response. To assess this, wild-type mice received bilateral ICV injections of 1mM (225μg/10μL dose) HTT-targeting monovalent siRNA or HTT-targeting blunt-di-siRNA^HTT^ **(Figure 6D)**. For each siRNA, two mice were injected side by side; one with the oligo resuspended in aCSF+, and the other with the oligo resuspended in 1XPBS buffer. The severity of acute neurotoxicity was similar between all four siRNAs injected 1mM in 1XPBS **(Figure 6E).** Formulating the siRNAs in aCSF+ significantly reduced the acute neurotoxicity and prevented any seizures from occurring **(Figure 6E)**. The blunt-ended di-siRNA was slightly less tolerated than the di-siRNA^HTT^ in aCSF+. This may be due to the blunt-ended molecule containing ten more phosphates than the di-siRNA^HTT^, allowing for more binding of divalent cations **(Figure 6D)**, as well as the higher avidity of divalent cations for double-stranded rather than single-stranded RNA.^21,51^ Further evaluation of the Ca^2+^ and Mg^2+^ concentration in aCSF buffer may be necessary when the total di-siRNA charge is changed, to completely mitigate all adverse events during recovery.

### Using aCSF+ reduces the acute neurotoxicity of CNS-administered ASOs

Fully PS ASOs induce severe short-term behavior phenotypes similar to siRNAs. The reduction in the PS context results in a decrease in the level of observed toxicity while sacrificing long-term stability.^19^ The most clinically advanced ASO GapmeRs utilize a mixed PO/PS backbone. We explored the potential of using the aCSF+ buffer on the safety profile of a full PS and a mixed PO/PS ASO in the CNS of wild-type mice **(Figure 7A)**. Tominersen^28^, one ASO tested here, is a published DNA-MOE GapmeR ASO that targets *Htt* to reduce total huntingtin RNA and protein **(Figure 7B)**. Substitution of the six PS backbones in the flank regions to PO significantly decreased the observable acute toxicity, similar to what is reported in a side-by-side comparison by Moazami et al.^19^ for C9orf72 targeting antisense. Using the same acute tolerability score described previously **(Figure 1C),** a comparable dose of 225μg/10μL of the full PS and mixed PS/PO ASOs was evaluated in wild-type mice under isoflurane **(Figure 7A)**. This dose of ASO has the same PO/PS molar equivalents as the dose used for the previously described siRNA safety evaluations. Mice injected ICV with fully PS-modified ASO showed severe neurotoxic phenotypes with observable seizures for extended periods of time, similar to siRNAs **(Figure 7C)**. The PS/PO mix backbone ASO was significantly better tolerated **(Figure 7D)** without detectable seizures by EEG/EMG **(Figure S7).** Instead, the mixed backbone ASO-injected mice consistently display hyperactivity during recovery, and some experienced ataxia. The time it took mice to wake-up and right themselves was less influenced by ASO-induced toxicity **(Figure S8)**, than what was seen with the di-siRNA studies **(Figure 1E, Figure S4)**.

**Figure 7.**
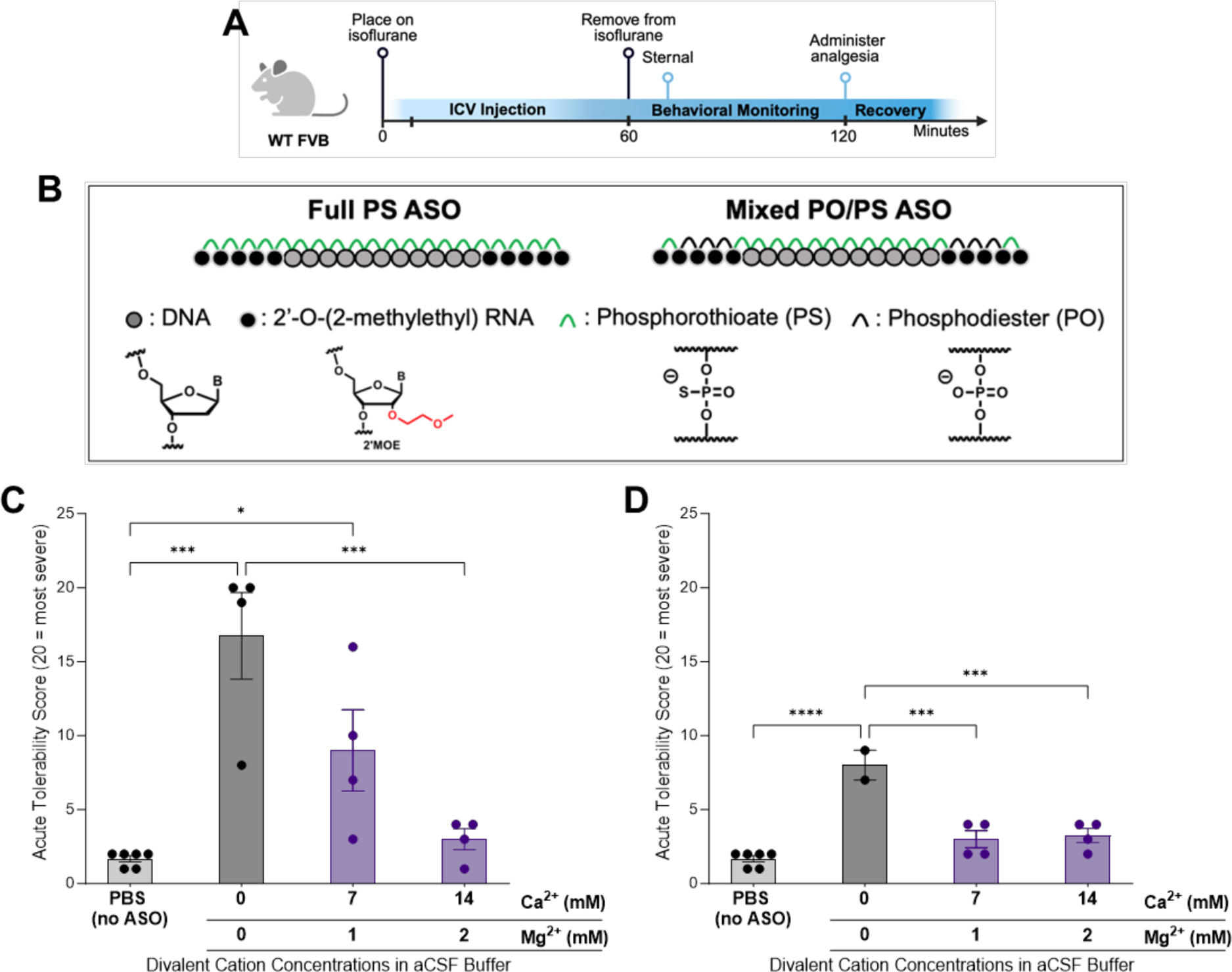
aCSF+ enables safe delivery for full PS and mixed PO/PS backbone ASOs. (A) Timeline showing the surgical procedure and recovery period for wild-type mice injected bilateral ICV with oligonucleotides under isoflurane anesthesia. (Created with BioRender.com) **(B)** The chemical structures of the full PS ASO and mixed PO/PS ASO (Tominersen^1^) used in this figure. **(C)** Acute tolerability scores following ICV injections of 225μg/10μL full PS ASO^HTT^ in 1XPBS or aCSF+ buffers. ICV injections of the full PS ASO produced significant acute neurotoxicity compared to 1XPBS controls (***p=<0.0001). Formulating ASO in two different aCSF+ buffers significantly improved the acute tolerability in a dose-dependent manner compared to 1XPBS controls (*p=0.0417 and ns). The 14:2 aCSF+ significantly improved the acute tolerability score compared to ASO in 1XPBS (***p=0.0008). Each data point represents one mouse (n=4-6) with SEM; data were analyzed using one-way ANOVA followed by Tukey’s post hoc test. **(D)** Acute tolerability scores following ICV injections of 225μg/10μL dose of mixed PO/PS ASO in 1XPBS or aCSF+ buffers. ICV injections of the mixed PO/PS ASO produced significant acute neurotoxicity compared to 1XPBS controls (****p=<0.0001). Formulating ASO in two different aCSF+ buffers significantly improved the acute tolerability (***p=0.0003 and ***p=0.0002). There was no difference in tolerability between the ASO in both aCSF+ buffers and the 1XPBS controls. Each data point represents one mouse (n=2-6) with SEM; data were analyzed using one-way ANOVA followed by Tukey’s post hoc test.

We next decided to evaluate the impact of the addition of Ca^2+^ and Mg^2+^ in aCSF with CSF-administration of ASOs. The duplexed (siRNA) and single-stranded (ASO) oligonucleotides differ profoundly in their biophysical properties.^52^ The ASO is amphiphilic in nature with hydrophobic bases and a negatively charged backbone. The duplexed part of the siRNA is rigid and negatively charged, with hydrophobic bases hidden inside the duplex. Thus, the optimal concentration of divalent might be different for two classes of oligonucleotides. For this reason, two formulations of aCSF+ were evaluated with 14:2 (14mM Ca^2+^ and 2mM Mg^2+^) and 7:1 (7mM Ca^2+^ and 1mM Mg^2+^) divalent cations added.

With the fully PS-modified ASO, both aCSF+ formulations increased the short-term tolerability of ASOs in a dose-dependent manner. The higher, 14:2 Ca^2+^ and Mg^2+^ formulation, as selected for the previously described di-siRNA studies, eliminated most acute adverse events during recovery **(Figure 7C).** For some animals, this formulation resulted in delayed wake-up times. With the 7:1 Ca^2+^ and Mg^2+^, some adverse events indicative of seizures were observed but were significantly reduced compared to the 1XPBS ASO delivery **(Figure 7C).** Thus, optimal ratios of the additional divalent cations in the formulation are likely different depending on the structure of the oligonucleotides.

For the mixed PO/PS backbone ASO, formulating in two different aCSF+ buffers for ICV injections significantly increased the acute tolerability in all mice, with no hyperactivity and ataxia phenotypes detected **(Figure 7D).** Similar to the full PS variants, longer recovery times were observed with the 14:2 Ca^2+^ and Mg^2+^ formulation, indicating that a non-bound excess of Ca^2+^ and Mg^2+^ might be problematic.

### The use of aCSF+ mitigates acute neurotoxicity of di-siRNA^HTT^ in Huntington’s disease mouse models

Finally, it was essential to confirm the acute safety of delivering di-siRNA^HTT^ **(Figure 8A)** in aCSF+ in neurodegenerative disease models under isoflurane anesthesia **(Figure 8B).** Before aCSF+ formulation, 1mM (225μg/10μL dose) di-siRNA^HTT^ delivered in 1XPBS to YAC128 HD **(Figure 8C)** and BAC-CAG HD **(Figure 8D)** transgenic mice elicited tonic-clonic seizures. With the aCSF+, 1mM di-siRNA^HTT^ was injected bilaterally into both lateral ventricles with no observable neurotoxicity in YAC128 HD mice **(Figure 8E)** and BAC-CAG HD mice **(Figure 8F).** Evaluating this delivery formulation in animal models with sporadic seizures may be beneficial to see if this level of divalent cations would enhance or dilute that phenotype. The mice were sacrificed one-month following injection, and HTT protein levels were measured by ProteinSimple Wes **(Figure 8G-J, Figure S9)** to confirm efficacy similar to what was seen in WT mice **(Figure 5)** and previously shown in HD models.^34,35,53^ Ensuring the safety of oligonucleotides in animal disease models should be an essential consideration before moving them into clinical trials.

**Figure 8.**
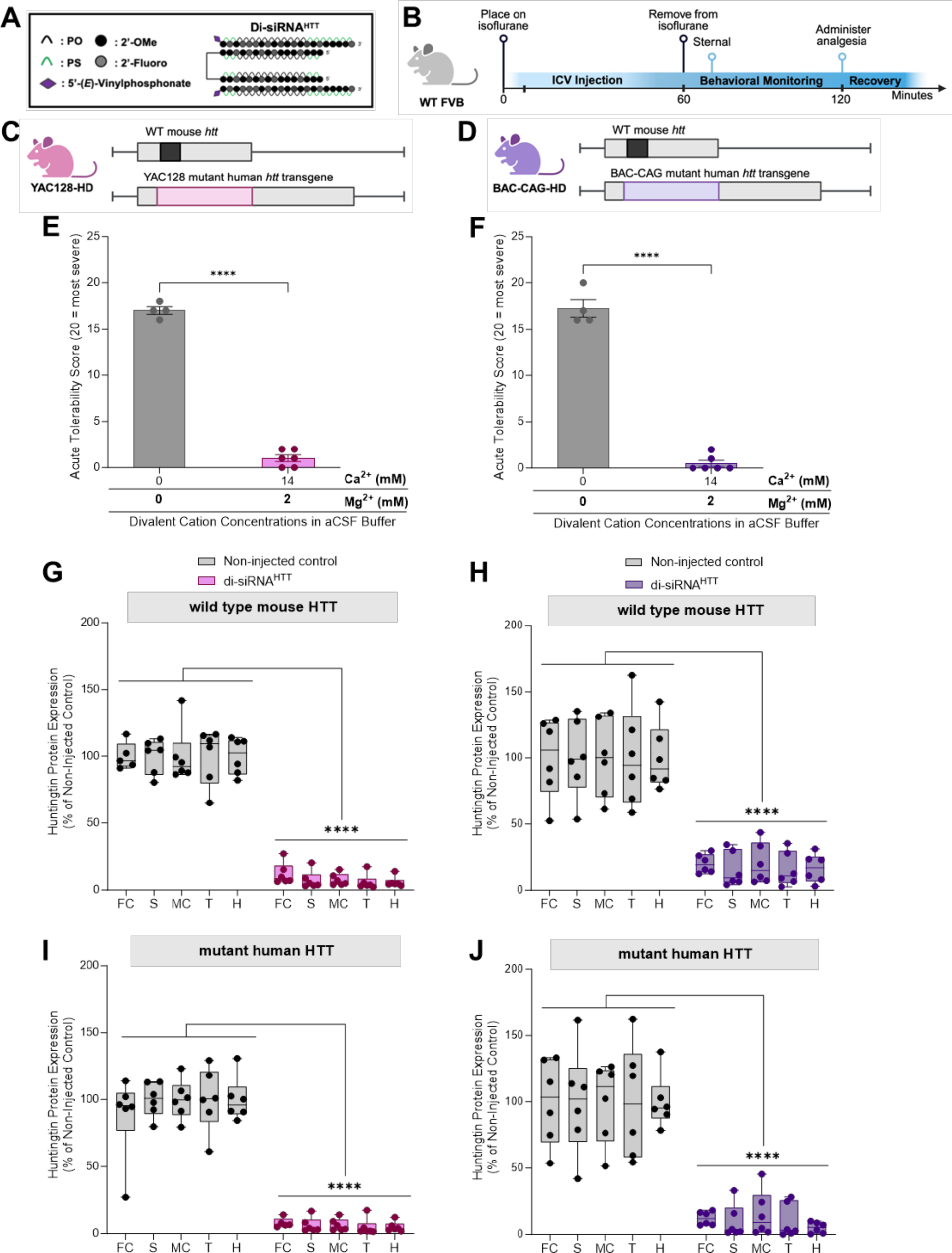
The use of aCSF+ mitigates acute neurotoxicity of di-siRNA^HTT^ in Huntington’s mouse models of disease. (A) The chemical structure of di-siRNA^HTT^ used throughout this figure. **(B)** Timeline showing the ICV injection procedure and recovery period for mice injected with oligonucleotides under isoflurane anesthesia. (Created with BioRender.com) **(C-D)** Schematics depicting the mouse models used in this figure, YAC128-HD^2^ and BAC-CAG-HD^3^. Both models carry two copies of wild-type mouse huntingtin (*Htt*) (black) and either a YAC (pink) or BAC (purple) transgenic insert containing human mutant *Htt.* (Created with BioRender.com) **(E, F)** ICV administration of 1mM (225μg/10μL total) di-siRNA^HTT^ formulated in 1XPBS elicited seizures in YAC128-HD and BACCAG-HD mouse models. **(E)** The acute tolerability was significantly increased in YAC128-HD mice following ICV administration of 1mM (225μg/10μL total) di-siRNA^HTT^ formulated in aCSF+. Each data point represents one mouse (n= 6). ****p=<0.0001; data were analyzed using an unpaired t-test. (**F)** The acute tolerability was significantly reduced in BACCAG-HD mice following ICV administration of 1mM di-siRNA^HTT^ formulated in aCSF+. Each data point represents one mouse (n= 6). ****p=<0.0001; data were analyzed using an unpaired t-test**. (G, H)** Wild-type mouse HTT and human mutant HTT **(I, J)** protein levels in di-siRNA^HTT^ treated mice measured by ProteinSimple Wes. One month following injections, there was a significant lowering of wild-type and mutant HTT protein across all brain regions measured versus the non-injected control groups in YAC128-HD **(G, I)** and BAC-CAG HD **(H, J)** mice. Each data point represents one mouse, and the error bars represent the mean values of SEM (n=6). ****p=<0.000; data were analyzed using two-way ANOVA with Tukey’s multiple comparisons test.

## Discussion

Oligonucleotides (ASOs and siRNAs) have become an increasingly popular therapeutic modality, with the CNS being one of the areas of intense preclinical and clinical development.^54,55^ Oligonucleotides do not cross the blood-brain barrier and must be delivered through direct cerebral spinal fluid (CSF) administration via intracerebroventricular (ICV) or intrathecally (IT) injections.^13^ While there are different types of toxicity associated with oligonucleotides ^47,48,50^, CSF administration will likely impose additional limitations. CSF is a tightly balanced fluid circulating throughout the brain, and its correct composition is essential for neuronal function and signal transduction. Calcium is one of the vital components of CSF that is directly involved in neuronal activity.^56^ Calcium and magnesium divalent cations play critical roles in neuronal functioning, and a reduction of either in the CSF and intracellular stores could lead to seizures.^57,58^ Although the mechanism remains unclear, there is evidence of hypocalcemia-induced seizures in humans with pre-existing endocrinological abnormalities.^59^ Thus, care must be taken to ensure that drugs administrated directly into the CSF have CSF-compatible formulations.

The molecular mechanisms underlying dose-limiting toxicities of oligonucleotides are diverse and not fully understood. Numerous clinical trials involving ASOs have been halted due to toxicity and an inability to reach effective levels, while others resulted in the development of disease-modifying drugs that change the lives of thousands of patients. Moreover, it is probable that various structural and chemical classes of oligonucleotides (single-stranded, double-stranded, lipophilic conjugates, divalent, etc.) may differ in their toxicity mechanisms. Clinical data on CSF-administered siRNAs is limited, with only one compound (ALN-APP)^11^ reaching clinical trials and demonstrating a significant reduction in APP cerebrospinal fluid biomarkers. The maximum tested dose in clinical trials was 75mg, constrained by reported toxicity observed in high-dose repetitive non-human primate studies (Nature Conference 2023).

Jia et al^15^ have recently reported a systematic evaluation of the acutely neurotoxic ASOs of different sequence and modification patterns. They demonstrate that the molecular mechanism driving neurotoxicity in toxic ASOs is related to the reduction in intracellular Ca^2+^ concentration in neurons, which can be mitigated by modulating Ca^2+^ channel activity. The authors explored the impact of preformulating ASOs with Ca^2+^ in vivo and observed a slight increase (∼10-15%) in observable short-term cumulative neurotoxicity. This finding contrasts with Moazami et al.^19^ and our own reports. Their study focused on neurotoxic ASOs at concentrations significantly lower (25 vs. 225μg), with observable neurotoxicity mostly limited to consciousness, motor function, and appearance, rather than profound seizures and death. It’s possible that other toxicity mechanisms, such as protein binding, may be contributing factors. Further systematic exploration of observable toxicity and mitigation strategies at a wide range of concentrations and formulations would be necessary to better understand the differences in reported experiments.

To avoid the complications associated with phosphate calcium salt precipitates, we used a HEPES-based^60^ aCSF buffer for all experiments. There is a limited number of pharmaceutical-grade buffers certified for CSF administration, with PBS being one of them. As buffering of nucleic acids is highly desired to ensure long-term stability and consistency, the use of calcium-containing formulations in the clinic would necessitate additional optimization and likely require the use of non-phosphate-based buffering solutions.

In rodents, long-term anesthesia, such as tribromoethanol (TBE, Avertin)^33^, was sufficient to mask the short-term neurotoxicity of high doses of siRNA. After an hour, mice recovered normally and demonstrated robust efficacy with no evidence of long-term toxicity. In our experiments, the optimized aCSF+ formulation effectively eliminated acute toxicity observed under reversible anesthesia for various di-siRNAs, including compounds targeting MSH3^61^, ApoE^34^, and others. For double-stranded RNAs, neurotoxicity appears more dependent on the scaffold than the sequence within the tested experimental conditions.

With ASOs, both sequence and modification patterns impact the degree of observable adverse events. Indeed, in this work, the HTT-targeting ASO GapmeR with a mixed versus fully phosphorothioated backbone exhibited profoundly different short-term toxicity. The fully PS-modified ASO induced seizure-like adverse events similar to siRNAs, while the mixed PO/PS backbone was better tolerated, with only hyperactivity and ataxia observed. For both ASOs, the use of aCSF+ reduced the short-term toxicity. For the injection dose of 225μg/10μL of (1mM) di-siRNA, a 14:2 (14mM Ca^2+^ and 2mM Mg^2+^) supplementation was fully effective in reducing observable short-term seizures. For 225μg/10μL of ASO (equivalent molar concentration of PO/PS backbones to the 225μg/10μL of (1mM) di-siRNA dose), aCSF+ with the same divalent cation content was effective in reducing observable short-term seizures, but the mice still experienced some other adverse events, including ataxia and extended wake-up times. These differences may be due to slight differences in the total charge of the molecule and should be evaluated further depending on the oligonucleotide.

The other interesting differences we observed between siRNAs and ASO is the senstitivty of the short-term toxicity to phosphorothioate content. PS backbone modifications enhance ASO toxicity, but the direct mechanism is not well defined.^62^ In the clinic, all GapmeR ASOs have a mixed PO/PS backbone where several PS linkages are substituted with PO to reduce the observable toxicity.^6,18,19^ Some studies have shown that PS-ASOs bind more frequently to nuclear proteins than PO-ASOs, which is associated with this toxicity.^63^ Consistent with published data, reduction in the PS context profoundly increased the short-term tolerability of the ASOs. With siRNAs, we have not seen such a profound impact correlated to the PS context. The toxicity of the siRNAs with 23% and 37% PS backbones was similar. Although the version of di-siRNA^HTT^ with no PS modifications (this siRNA is functionally inactive) was slightly better tolerated than the 26mM-PS modified version of di-siRNA^HTT^ in 1XPBS, the same concentration of divalent cations in aCSF was required to mitigate seizures and adverse events for both completely.

A different formulation may be necessary for some ASOs and siRNAs due to the profoundly different biophysical properties of the two oligonucleotide classes. ASOs are single-stranded and thus amphiphilic in nature, while siRNAs are duplex, rigid, and negatively charged. Double-stranded oligonucleotides also have a higher binding capacity for binding divalent cations compared to single-stranded oligonucleotides.^51^ Another potential difference is the sensitivity of the observed effects to sequences. With single-stranded ASOs, sequence profoundly affects the self-structure and, thus, the degree of amphiphilicity, resulting in an impact on divalent cation binding avidity. Additionally, self-folding can create dimeric or aptameric-like effects, where the exact mode of interaction with ions and cells is challenging to predict. This is likely why this class of toxicity is more affected by sequence and chemical modification patterns in ASOs but not in siRNAs.^64–66^

Overall, our studies provide *in vivo* validation of oligonucleotide-induced neurotoxicity following ICV injections in mice and prevention methods. The use of EEG/EMG in awake animals sets our studies apart from others in the field, as it provides quantifiable measurements of oligonucleotide-induced seizures. Divalent cations bind preferentially to the phosphate backbone of oligonucleotides, which translates to an imbalance in the ion homeostasis of the CSF following ICV injections in *vivo*. This imbalance can be prevented when oligonucleotides are delivered with specific divalent cations in aCSF buffer. We observed that formulation in aCSF+ has no impact on functional silencing or other measurable phenotypes. Thus, in preclinical models, divalent cation-enriched buffers will simplify experiments and allow higher doses of oligonucleotides to be administered, which may enable longer duration and less frequent administration requirements. The translation to clinic is much more complicated, and detailed studies on the toxicity and tolerability of divalent cation enriched formulation need to be performed in large animal models and clinics. This area of research deserves further exploration to understand the deeper mechanisms of ionic imbalance in the CSF and the long-term implications on brain functioning and neuronal survival.

## MATERIALS AND METHODS

All of the studies in this manuscript were conducted in compliance with the Institutional Care and Use Committee (IACUC) guidelines of the University of Massachusetts Chan Medical School (docket #202100018 and A2574) and the University of California Davis (docke t#22308).

### Statistics and Reproducibility

Data analyses were performed using GraphPad Prism 9.0 software (GraphPad Software Inc.). Data were analyzed using a one-way or two-way ANOVA test with Bonferroni’s or Tukey’s test for multiple comparisons as specified in the figure legends. Differences in all data sets were considered significant at p < 0.05.

### Oligonucleotide Synthesis

Oligonucleotides were synthesized by phosphoramidite solid-phase synthesis on an AKTA Oligopilot 100 (Cytiva, Marlborough, MA) with standard protocols. 5’-(E)-Vinyl tetraphosphonate (pivaloyloxymethyl) 2’-O-methyl-uridine 3’-CE phosphoramidite was used for the addition of 5’-Vinyl Phosphonate, 2ʹ-F, 2ʹ-OMe phosphoramidites with standard protecting groups were used to make the modified siRNAs (Hongene Biotech, Union City, CA). Phosphoramidites were dissolved at 0.1 M in anhydrous acetonitrile (ACN), with added anhydrous 15% dimethylformamide in the case of 2’-OMe-Uridine amidite. 5-(Benzylthio)-1H-tetrazole (BTT) was used as the activator at 0.25 M. Coupling times were 4 minutes. Detritylations were performed using 3% trichloroacetic acid in dichloromethane. Capping reagents used were CAP A (20% N-methylimidazole in ACN) and CAP B (20% acetic anhydride and 30% 2,6-lutidine in ACN). Phosphite oxidation to convert to phosphate or phosphorothioate was performed with 0.05 M iodine in pyridine-H2O (9:1, v/v) or 0.1 M solution of 3-[(dimethylaminomethylene)amino]-3H-1,2,4-dithiazole-5-thione (DDTT) in pyridine for 3 minutes, respectively. All reagents were purchased from Chemgenes, Wilmington, MA. Guide oligonucleotides were synthesized on 500 Å long-chain alkyl amine (LCAA) controlled pore glass (CPG) functionalized with Unylinker terminus (Chemgenes). Divalent oligonucleotides (DIO) were synthesized on custom solid support prepared in-house.^12^

### Deprotection and purification of oligonucleotides

Vinyl-phosphonate containing oligonucleotides were cleaved and deprotected with 3% diethylamine in ammonium hydroxide, for 20 hours at 35℃ with agitation. DIO oligonucleotides were cleaved and deprotected with 1:1 ammonium hydroxide and aqueous monomethylamine, for 2 hours at 25℃ with agitation. The controlled pore glass was subsequently filtered and rinsed with 30mL of 5% ACN in water, solutions were evaporated overnight using a centrifugal vacuum concentrator. Purifications were performed on an Agilent 1290 Infinity II HPLC system (Agilent Technologies, Lexington, MA), using Source 15Q ion exchange resin (Cytiva). The loading solution used was 20 mM sodium acetate in 10% ACN in water, and elution solution was the loading solution with 1M sodium bromide. A linear gradient was used from 30 to 70% in 40 min at 50°C. Peaks were monitored at 260nm. Pure fractions were combined and desalted by size exclusion with HPLC Honeywell water AH365 using Sephadex G-25 (Cytiva). The purity and identity of fractions and pure oligonucleotides were confirmed by IP-RP LC/MS (Agilent 6530 Accurate-mass Q-TOF LC/MS).

### Ion Chromatography

Small aliquots (0.2mL) of the di-siRNAs (4mM) in sodium form dissolved in water were mixed 1:1 with solutions of 200mM Ca^2+^, 200mM Mg^2+^, or 200mM Ca^2+^/Mg^2+^, forcing exchange of the divalent cations to the bound monovalent sodium cations. Removal of the excess cations in solution not bound to the siRNAs was done with 3K centrifugal filter units (Amicon® Ultra-4). siRNAs were diluted to a final volume of 4mL with nuclease-free water and transferred to the Amicon tubes. Tubes were then centrifuged 40min at 5000 rpm (Wash). The wash procedure was repeated a total of three times, where each time the holdup volume (∼0.1mL) was diluted to a final of 4mL to start the next wash. The osmolality of the solution was determined after each wash to track the removal process. Osmolality was measured on a FreezePoint 6000 Osmometer (ELITechGroup, Logan, UT).

Ion chromatography was performed on a Metrohm Compact IC Flex, equipped with an autosampler and conductivity detector (Metrohm, Riverview, FL). 20uL of di-siRNA was loaded into a Metrosep C4 4×150mm column. A 12-minute isocratic eluent of 3.5mM Oxalic Acid solution was used to separate the cations. A calibration curve in the linear range of 1 – 75ppm of standards mix (Li^+^, Na^+^, NH_4_^+^, K^+^, Mg^2+^, and Ca^2+^) was used to calculate the concentration of the studied cations in the samples.

### Mouse Intracranial Injection Surgeries

WT FVB mice were purchased from The Jackson Laboratories. YAC128-HD^67^ and BAC-CAG-HD^68^ mice were maintained in a colony as heterozygotes on an FVB background. All animals were maintained and used according to the Institutional Animal Care and Use Committee guidelines of the University of Massachusetts Chan Medical School (docket #20210018). Briefly, mice were housed with a maximum of 5 per cage in a pathogen-free facility under standard conditions with access to food, water, and enrichment *ad libitum.* Mice were injected bilaterally directly into the ICV at eight weeks of age. The mice were anesthetized under constant isoflurane flow and placed into a small animal stereotaxic apparatus (ASI Instruments and WPI Instruments).

Microinjector pumps were fastened to the stereotaxic and set to deliver 5.0µL of the siRNA at a rate of 500 nL/min. The surgical position was set using the bregma as the zero position for guidance, then measured +/-0.8 mm medial-lateral and -0.2 mm anterior-posterior (AP), and finally lowered -2.5mm dorsal-ventral (DV) into the ICV. The injection is started one minute after the needle is inserted, and the needle is removed one minute after the injection is completed. Once the injection was complete, the mice were placed on a heating pad until fully awake and returned to their primary housing facility. Each animal received 0.1cc/10g Ketofen subcutaneously once fully awake. All mice were housed individually post-op to allow healing and avoid losing stitches.

### Mouse Tissue Collection

The experimental endpoint for mRNA and protein analysis was one or two months post-ICV injections. At the determined time point, mice were deeply anesthetized with tribromoethanol and perfused intracardially with 20mL cold 1X PBS buffer. The brains were cut in half; half was post-fixed in 10% formalin for 24 hours and switched to 1X PBS at 4°C for long-term storage. The other half was immediately sliced into 1.0mm sections using a brain matrix (Kent Scientific Corporation, RBMA-200C). Brain sections floating in cold 1X PBS were punched using 1.5mm sterile biopsy punches. The brain punches were submerged in RNAlater or flash-frozen on dry ice and stored at -80°C for mRNA and protein analysis.

### Electrophysiology in Mice: Surgery

Mice were anesthetized with ketamine/xylazine [100 and 10 mg/kg, respectively, intraperitoneally (IP)] and then placed in a stereotaxic apparatus. Mice were implanted with a bi-lateral guide cannula (PlasticsOne, #C235GS) in the lateral ventricles (coordinates from Bregma are 0.0 mm Antero-posterior, ±1.0 mm Lateral, −2.3 mm Dorso-ventral, as per the mouse atlas of Paxinos and Franklin 2001). ^69,7071,72^ The guide cannula is closed using a dummy (PlasticsOne, #C235DCS) and secured using a dust cap (PlasticsOne, #303DC). The mice were also implanted with two EEG screw electrodes, one parietal (mid-distance between bregma and lambda and 1 mm lateral from the mid-line) and one on the cerebellum (6.5 mm caudal, 0 mm lateral from bregma, reference electrode) electrodes (Pinnacle Technology Inc., Catalog #8403) and two flexible electromyogram (EMG) wire electrodes (in the neck muscles; Plastics One, catalog #E363/76/SPC), previously soldered to a 6-pin connector (Heilind Electronics, catalog #853-43-006-10-001000). The whole assembly is secured to the skull with dental cement. After completing the surgery, mice were kept in a warm environment until normal activity was resumed, as previously described. ^36–40^

### Electrophysiology in Mice: Surgery: *Sleep-Wake Recording*

Following the surgery, the mice are housed individually in transparent barrels in an insulated sound-proofed recording chamber maintained at an ambient temperature of 22 ± 1 ◦C and on a 12 h light/dark cycle (lights-on at 07:00, Zeitgeber time: ZT0) with food and water available *ad libitum*. After a minimum of 9 days for post-surgical recovery, mice are connected to flexible recording cables and habituated to the recording conditions for five days before starting polygraphic recording. The cortical EEG and EMG signals are amplified (A-M System 3500, United States) and digitalized with a resolution of 256 Hz using Vital Recorder (Kissei, Japan). Video recording is synchronized to the EEG/EMG recording. Mice are recorded for a 24 h baseline period. Then, the mice received bi-lateral ICV injection (5 μl/side, 0.5 μl/min, 10 min separating the first side injection from the second side injection). Briefly, awake mice, without disconnecting the EEG/EMG cable, were gently maintained still to remove the dust cap and the dummy from the guide cannula and insert the injector system (PlasticsOne, #C235IS) previously connected to a 40cm long tubing connector (PlasticsOne, #C232C), itself connected to two Hamilton syringes (5μl each). The mice were then placed back in the home cage and freely moved during the entire injection period. Ten minutes after completion of the bilateral ICV injection, the mice were again gently maintained still in order to remove the injector system and insert the dummy and the dust cap back in. EEG/EMG signals were recorded for 24 hrs following the end of the ICV injection. The mice were then deeply anesthetized (ketamine/xylazine 200 and 20 mg/kg, respectively, IP) and injected ICV with a blue dye (0.5-1 μl), and the brain was dissected to control for guide cannula correct placement in the lateral ventricles **(Figure S2).** Mice with incorrect guide cannula placement were removed from the study.

### Electrophysiology in Mice: *EEG/EMG Analysis*

Using SleepSign for Animal (Kissei, Japan) assisted by spectral analysis using fast Fourier transform (FFT), polygraphic records are visually inspected for high amplitude EEG/EMG waves characteristic of seizures. Seizures were defined as periods with high frequency and high amplitude with a minimum duration of 5 s. Seizures were identified visually from EEG/EMG signals, and videos were checked to determine if there was a behavioral accompaniment.

### Htt mRNA Silencing Quantification

Brain tissue punches were incubated in RNAlater overnight and then stored at -80°C. The punches were lysed in 600μL Quantigene homogenizing solution (ThermoFisher Cat#QS0518) with 6μL 20mg/mL proteinase K solution (ThermoFisher cat#AM2546) with sterile beads on a tissue lyser (Quantigene cat#9003240). The lysates were spun down (@2,000xg) and incubated at 55°C for 30 minutes. The lysates were transferred to deep well plates and stored at -80°C. On the day of analysis. Brain lysates were incubated at 55°C until thawed (approximately 30 minutes). The Quantigene manufacturer’s protocol was followed for the Singleplex Gene Expression Assay (ThermoFIsher cat#QS0016), as previously published.^35^ Wild-type mouse *Htt* mRNA (probe SB-14150) and human *HTT* mRNA (probe SB-50339) levels were measured and normalized to mouse HPRT mRNA (probe SB-15463). The background signal was subtracted, and all reads were normalized to non-injected controls.

### HTT Protein Silencing Quantification by ProteinSimple (Wes)

Frozen tissue punches were homogenized on ice in 75 µl 10mM HEPES pH7.2, 250mM sucrose, 1mM EDTA + protease inhibitor tablet (Roche) + 1mM NaF + 1mM Na3VO4, sonicated for 10 seconds and protein concentration determined by Bradford assay (BioRad). The standard settings for the 66-440 kDa separation modules were used with 0.2 mg/ml of each lysate and anti-HTT (Ab1^71^, 1:50) plus anti-Vinculin (Sigma, 1:2000) antibodies. The peak area was determined using Compass for SW software (ProteinSimple) and dropped line-fitted peaks.

### Divalent Cation Supplementation in aCSF Buffer

Stock solutions of artificial CSF (aCSF) were prepped as follows: 1M CaCl^2^, 1M MgCl^2^, 1M NaCl, 1M KCl^2^, and 100mM D-+-glucose CaCl^2^, all in sterile water. The stock aCSF diluent was prepped with the following final concentrations of components: 137mM NaCl, 5mM KCl2, and 20mM D-+-glucose in an 8mM HEPES buffer base. To prepare aCSF buffers with varying ratios of divalent cations, serial dilutions of CaCl^2^ and MgCl^2^ in the stock aCSF buffer were made. Finally, CaCl^2^ and MgCl^2^ containing aCSF stocks were combined to make buffers containing various ratios of divalent cations. After synthesis, the siRNAs resuspended in water were duplexed and dried in a Speedvac overnight. The resulting pellets were resuspended in the necessary aCSF buffer for the injected dose.

### Duplexing of siRNAs for Ion Supplementation in aCSF Buffer

Single strands of siRNAs were completely dried down using a SpeedVAC. Once dry, single strands were resuspended in pre-made aCSF containing Ca^2+^/Mg^2+^. Resuspended single-strand concentrations were measured by nanodrop, and the A260 value was used to calculate. Antisense strands were duplexed to dio-sense strands at a 2:1 ratio by heating to 95°C for 5 minutes and cooling at room temperature. Duplexes were verified by running on a 20% TBE agarose gel at 180V for 1 hour. Duplex concentrations were measured by nanodrop, and the A260 was used to calculate them. Depending on the desired dose, aCSF was added to dilute as necessary.

## Acknowledgments

The authors acknowledge Metrohm (Riverview, FL) for providing the ion chromatography equipment for the *in vitro* analysis of oligonucleotides (Figure 3). The authors thank Vicky Benoit, Kathryn Chase, and Lori Kennington for their assistance in breeding and genotyping mice.

## Authorship Confirmation/Contribution Statement

Conceptualization was performed by R.M., D.E., J.A., A.K., and N.A. Methodology was designed by R.M., D.E., J.A., J.W., A.K., and N.A. Investigation was performed by R.M., J.P., A.B., E.S., N.M., B.B., N.Y., and K.Y. Funding acquisition was done by C.A., M.D., A.K., and N.A. Project administration and supervision were performed by A.K., and N.A. Writing the original draft was completed by R.M., A.K., and N.A. Reviewing and editing the original draft was completed by R.M., D.E., J.W., M.D., A.K., and N.A.

