## Supplemental Materials for "Preventing acute neurotoxicity of CNS therapeutic oligonucleotides with the addition of Ca^2+^ and Mg^2+^ in the formulation"

**Table S1.** Oligonucleotide sequence and chemical modification patterns in this manuscript.

| **Oligonucleotide** | **Strand** | **Sequence (5'-3')** |
|---|---|---|
| di-siRNA$^{HTT}$ | Antisense | (VP)mU#fU#mA.fA.fU.fC.mU.fC.mU.fU.mU.fA.mC.fU#mG#fA#mU#mA#mU#mU#fU |
| di-siRNA$^{HTT}$ | Sense | mU#mC#mA.fG.mU.mA.mA.fA.mG.fA.mG.fA.mU.fU#mA#mA-DIO |
| di-siRNA$^{HTT}$ (NPS) | Antisense | (VP)mU.fU.mA.fA.fU.fC.mU.fC.mU.fU.mU.fA.mC.fU.mG.fA.mU.mA.mU.mU.fU |
| di-siRNA$^{HTT}$ (NPS) | Sense | mU.mC.mA.fG.mU.fA.mA.fA.mG.fA.mG.mA.mU.fU.mA.mA-DIO |
| Blunt- di-siRNA$^{HTT}$ | Antisense | (VP)mU#fU#mA.fA.fU.fC.mU.fC.mU.fU.mU.fA.mC.fU.mG.fA.mU.mA.mU#mU#fU |
| Blunt- di-siRNA$^{HTT}$ | Sense | mA#mA#mA.mU.mA.mU.mC.mA.fG.mU.fA.mA.fA.mG.fA.mG.mA.mU.fU#mA#mA-DIO |
| Monovalent siRNA | Antisense | (VP)mU#fU#mA.fA.fU.fC.mU.fC.mU.fU.mU.fA.mC.fU#mG#fA#mU#mA#mU#mU#fU |
| Monovalent siRNA | Sense | mU#mC#mA.fG.mU.mA.mA.fA.mG.fA.mG.fA.mU.fU#mA#mA |
| Full PS ASO | Antisense | eC#eU#eC#eA#eG#dT#dA#dA#d5C#dA#dT#dT#dG#dA#d5C#eA#eC#eC#eA#eC |
| Mixed PO/PS ASO | Antisense | eC#eU.eC.eA.eG#dT#dA#dA#d5C#dA#dT#dT#dG#dA#d5C#eA.eC.eC.eA#eC |

Chemical modifications are designated as follows: "." – phosphodiester bond, "#" – phosphorothioate bond, "m" – 2'-O-Methyl, "f" – 2'-Fluoro, "e" -2'-O-Methoxyethyl, "d" DNA, "DIO" – Glycerol-tetraethyleneglycol linker, "(VP)"- 5'-(E)-Vinylphosphonate.

**Table S2.** Average acute tolerability scores for all mice throughout the manuscript.

| Oligonucleotide | Dose (nmol) | Dose (μg) | Ca (mM) | Mg (mM) | Average Tolerability Score | Group Size (n=) | Mouse Genotype | Figure |
|---|---|---|---|---|---|---|---|---|
| 1X PBS Control | 0 | 0 | 0 | 0 | 1.67 | 6 | WT FVB | 1D, 6C |
| Di-siRNA$^{HTT}$ (P3, high PS) | 1.25 | 28.125 | 0 | 0 | 2.00 | 4 | WT FVB | 1D |
| Di-siRNA$^{HTT}$ (P3, high PS) | 2.5 | 56.25 | 0 | 0 | 2.50 | 4 | WT FVB | 1D |
| Di-siRNA$^{HTT}$ (P3, high PS) | 5 | 112.5 | 0 | 0 | 8.40 | 5 | WT FVB | 1D |
| Di-siRNA$^{HTT}$ (P3, high PS) | 10 | 225 | 0 | 0 | 14.2 | 5 | WT FVB | 1D |
| Di-siRNA$^{HTT}$ (P3, high PS) | 10 | 225 | 0 | 0 | 19.5 | 4 | WT FVB | 4C, 4D, 4E, 7C |
| Di-siRNA$^{HTT}$ (P3, high PS) | 10 | 225 | 4 | 0 | 11.75 | 4 | WT FVB | 4C |
| Di-siRNA$^{HTT}$ (P3, high PS) | 10 | 225 | 8 | 0 | 6.75 | 4 | WT FVB | 4C |
| Di-siRNA$^{HTT}$ (P3, high PS) | 10 | 225 | 12 | 0 | 3.25 | 4 | WT FVB | 4C |
| Di-siRNA$^{HTT}$ (P3, high PS) | 10 | 225 | 16 | 0 | 1.5 | 5 | WT FVB | 4C, 4E, 6C |
| Di-siRNA$^{HTT}$ (P3, high PS) | 10 | 225 | 32 | 0 | 1 | 4 | WT FVB | 4C |
| Di-siRNA$^{HTT}$ (P3, high PS) | 10 | 225 | 0 | 4 | 9.8 | 5 | WT FVB | 4D |
| Di-siRNA$^{HTT}$ (P3, high PS) | 10 | 225 | 0 | 8 | 8.5 | 4 | WT FVB | 4D |
| Di-siRNA$^{HTT}$ (P3, high PS) | 10 | 225 | 0 | 12 | 8.5 | 4 | WT FVB | 4D |
| Di-siRNA$^{HTT}$ (P3, high PS) | 10 | 225 | 0 | 16 | 2.75 | 4 | WT FVB | 4D |
| Di-siRNA$^{HTT}$ (P3, high PS) | 10 | 225 | 0 | 32 | 4 | 4 | WT FVB | 4D |
| Di-siRNA$^{HTT}$ (P3, high PS) | 10 | 225 | 15 | 1 | 0.5 | 4 | WT FVB | 4E |
| Di-siRNA$^{HTT}$ (P3, high PS) | 10 | 225 | 14 | 2 | 0.6 | 5 | WT FVB | 4E, 6C, 7C |
| Di-siRNA$^{HTT}$ (P3, high PS) | 10 | 225 | 13 | 3 | 1.5 | 4 | WT FVB | 4E |
| Di-siRNA$^{HTT}$ (P3, high PS) | 10 | 225 | 12 | 4 | 1.25 | 4 | WT FVB | 4E |
| Di-siRNA$^{HTT}$ (P3, high PS) | 10 | 225 | 11 | 5 | 4.75 | 4 | WT FVB | 4E |
| Di-siRNA$^{HTT}$ (P3, high PS) | 10 | 225 | 10 | 6 | 5.75 | 4 | WT FVB | 4E |
| Di-siRNA$^{HTT}$ (P3, high PS) | 20 | 450 | 16 | 0 | 12 | 2 | WT FVB | 6C |
| Di-siRNA$^{HTT}$ (P3, high PS) | 20 | 450 | 32 | 0 | 8.25 | 4 | WT FVB | 6C |
| Di-siRNA$^{HTT}$ (P3, high PS) | 20 | 450 | 28 | 4 | 1.25 | 4 | WT FVB | 6C |
| Di-siRNA$^{HTT}$ (P3, no PS) | 10 | 225 | 0 | 0 | 9.5 | 4 | WT FVB | 6E |
| Di-siRNA$^{HTT}$ (P3, no PS) | 10 | 225 | 14 | 2 | 1.25 | 4 | WT FVB | 6E |
| monovalent siRNA$^{HTT}$ | 10 | 225 | 0 | 0 | 13 | 4 | WT FVB | 6E |
| monovalent siRNA$^{HTT}$ | 10 | 225 | 14 | 2 | 1.75 | 4 | WT FVB | 6E |
| blunt-ended siRNA$^{HTT}$ | 10 | 225 | 0 | 0 | 16 | 2 | WT FVB | 6E |
| blunt-ended siRNA$^{HTT}$ | 10 | 225 | 14 | 2 | 2 | 4 | WT FVB | 6E |
| Full PS ASO | 10 | 225 | 0 | 0 | 16.75 | 2 | WT FVB | 7C |
| Full PS ASO | 10 | 225 | 14 | 2 | 9 | 4 | WT FVB | 7C |
| Full PS ASO | 10 | 225 | 7 | 1 | 3 | 4 | WT FVB | 7C |
| Mixed PO/PS ASO | 10 | 225 | 0 | 0 | 8 | 2 | WT FVB | 7D |
| Mixed PO/PS ASO | 10 | 225 | 14 | 2 | 3 | 4 | WT FVB | 7D |
| Mixed PO/PS ASO | 10 | 225 | 7 | 1 | 3.25 | 4 | WT FVB | 7D |
| Di-siRNA$^{HTT}$ (P3, high PS) | 10 | 225 | 0 | 0 | 17 | 4 | YAC128-HD | 8E |
| Di-siRNA$^{HTT}$ (P3, high PS) | 10 | 225 | 14 | 2 | 1 | 6 | YAC128-HD | 8E |
| Di-siRNA$^{HTT}$ (P3, high PS) | 10 | 225 | 0 | 0 | 17.25 | 4 | BAC-CAG HD | 8F |
| Di-siRNA$^{HTT}$ (P3, high PS) | 10 | 225 | 14 | 2 | 0.5 | 6 | BAC-CAG HD | 8F |

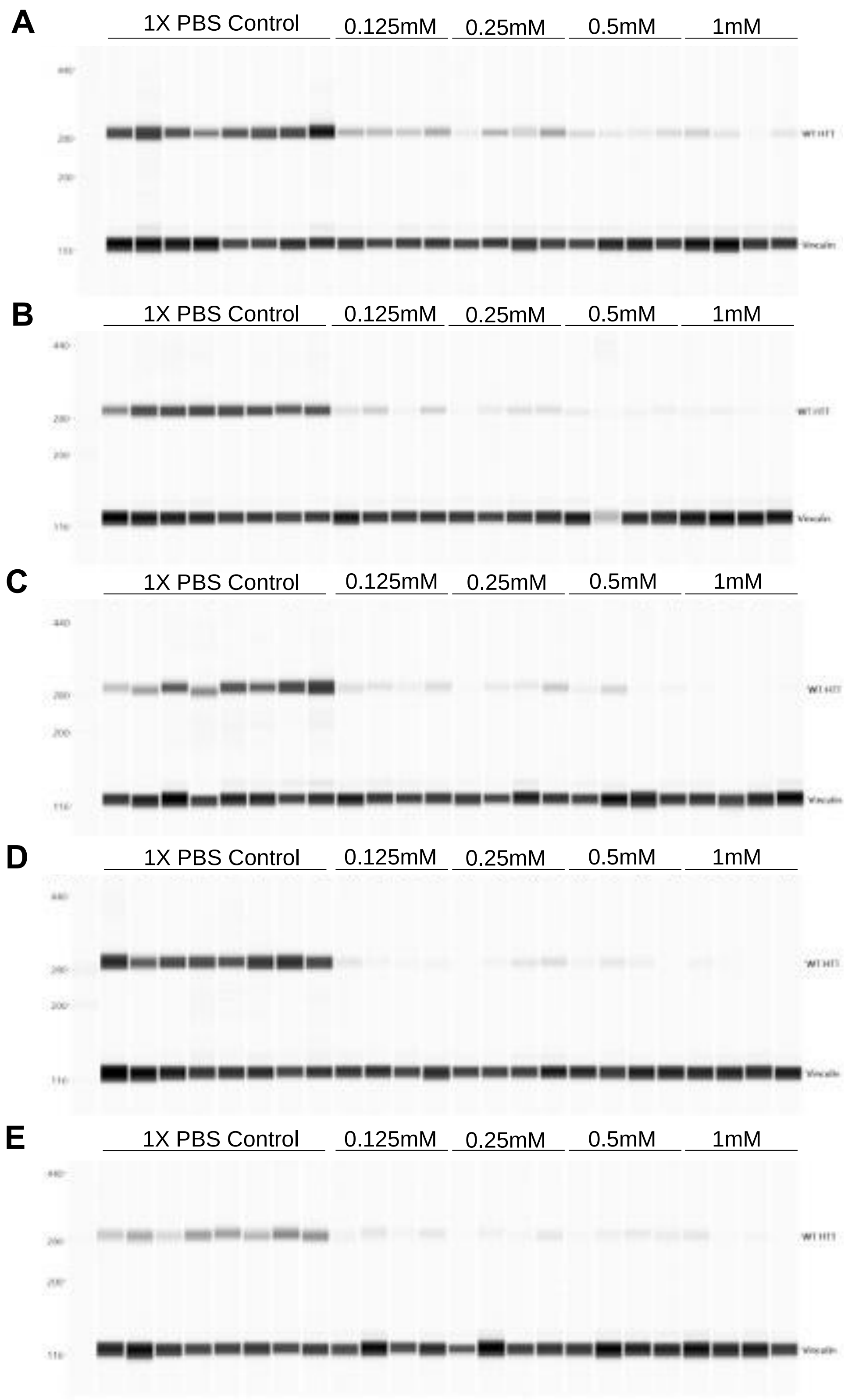
A
1X PBS Control
0.125mM
0.25mM
0.5mM
1mM
B
1X PBS Control
0.125mM
0.25mM
0.5mM
1mM
C
1X PBS Control
0.125mM
0.25mM
0.5mM
1mM
D
1X PBS Control
0.125mM
0.25mM
0.5mM
1mM
E
1X PBS Control
0.125mM
0.25mM
0.5mM
1mM

**Figure S1.** Raw western blot (ProteinSimple) of wild-type FVB/NJ mice treated with 0.25mM, 0.5mM, 1mM (~56μg, 112.5μg, 225μg/10μL total) di-siRNA$^{HTT}$ or 1XPBS control (Figure 1F). The levels of wild-type mouse HTT protein was evaluated in the frontal cortex **(A)**, striatum **(B)**, medial cortex **(C)**, thalamus **(D)**, and hippocampus **(E)** (n=4).

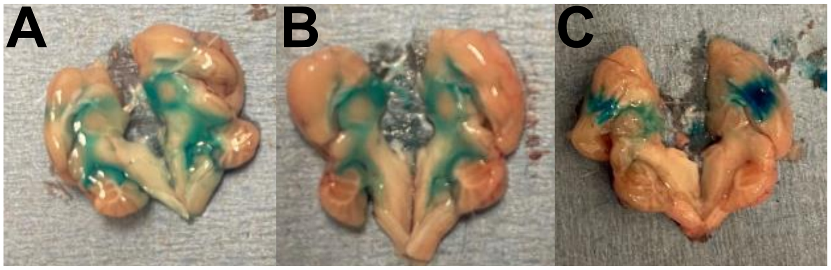


**Figure S2.** Confirmation of the guide cannula placement in the lateral ventricles for EEG/EMG studies. **(A, B)** At the end of the EEG/EMG recording, mice were deeply anesthetized (ketamine/xylazine, 200 and 20 mg/kg, respectively, intraperitoneally, IP), and blue dye (0.5μl) was injected ICV bilaterally. Mouse brains were extracted and sectioned / side (antero-posterior axis, mid-line). The blue dye is distributed evenly throughout the CSF stores, confirming the accurate placement of the guide cannula in the ventricles. **(C)** The blue dye is pooled in the brain tissue, specifically in the cortex, indicating that the guide cannula was positioned dorsal from the lateral ventricles in the cortex. This confirms the inaccurate or missed placement of the guide cannula. These animals are excluded from the study groups.

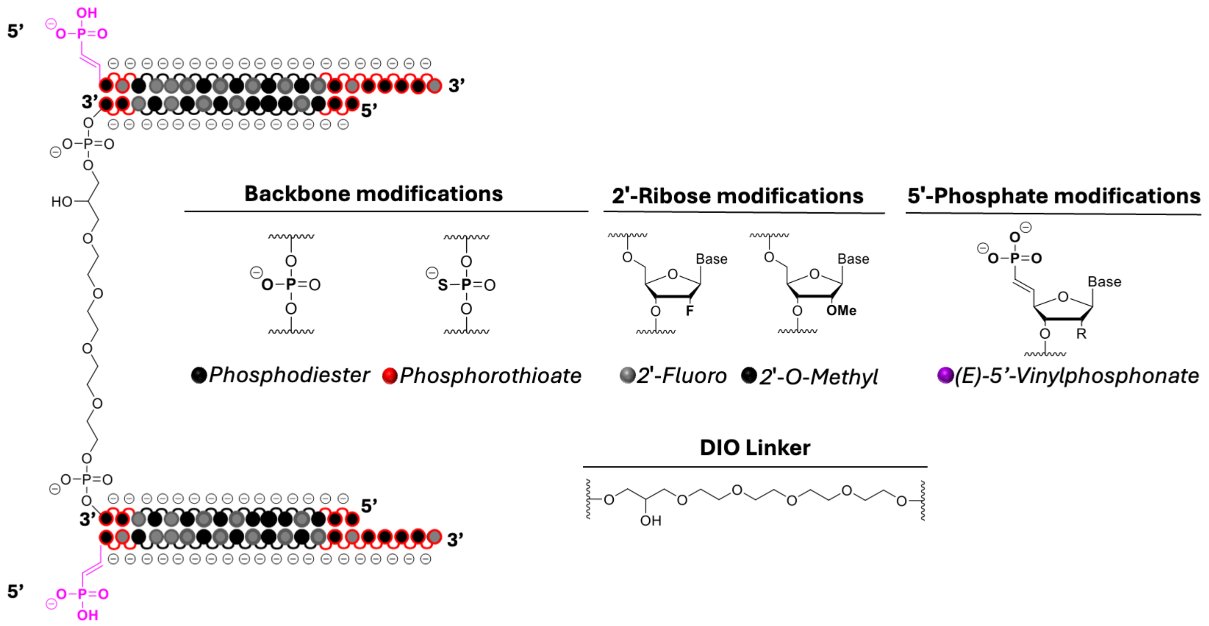


**Figure S3.** Schematic of the fully chemically modified di-siRNA$^{HTT}$ depicting the backbone charges and placement of binding opportunities. The negative charges can potentially bind a 1:1 ratio of up to 74 cations to maintain the charge neutrality of the solution.

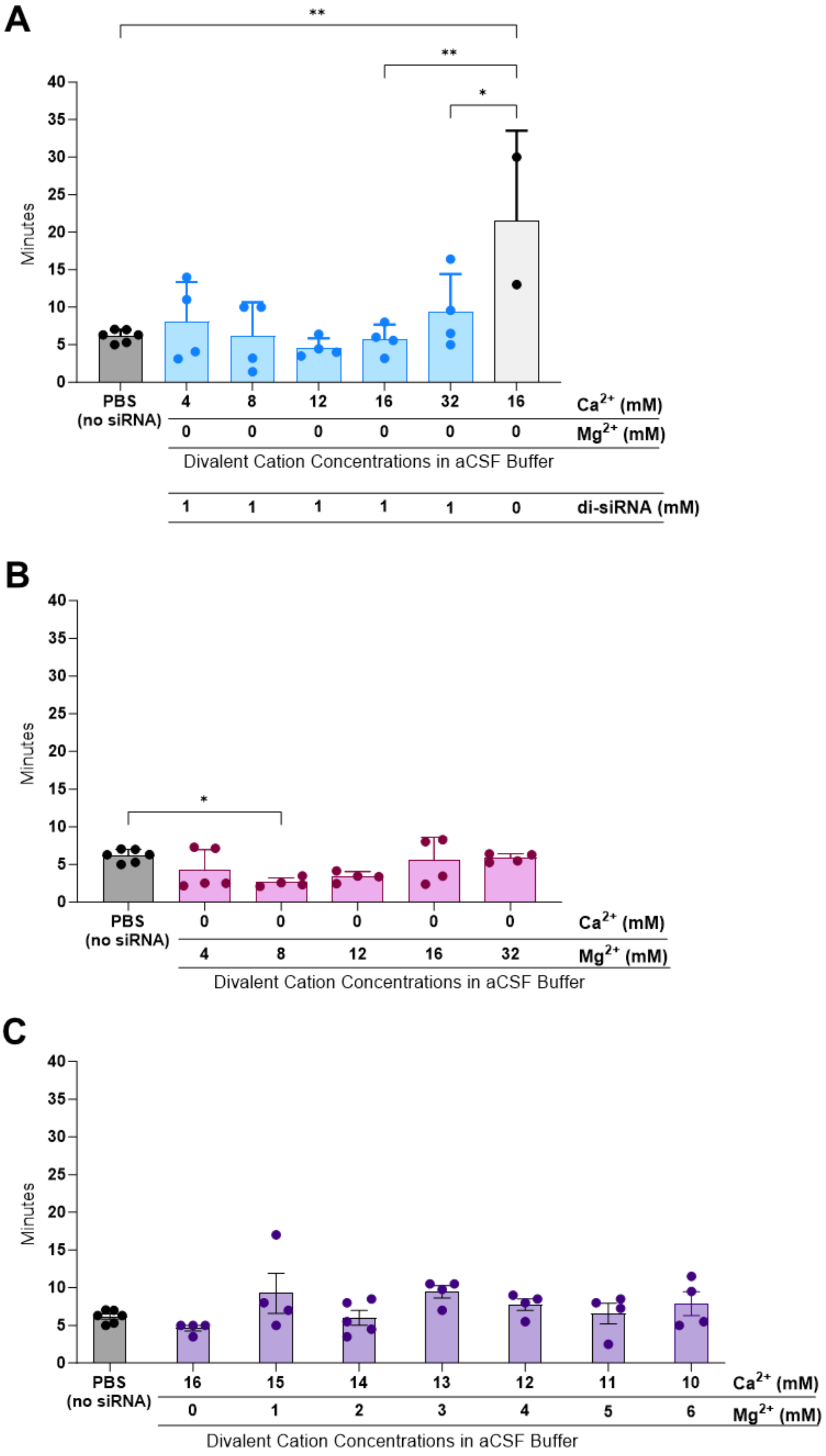
A
**
**
*
Minutes
40
35
30
25
20
15
10
5
0
PBS
(no siRNA)
4 8 12 16 32 16 Ca2+ (mM)
0 0 0 0 0 0 Mg2+ (mM)
Divalent Cation Concentrations in aCSF Buffer
1 1 1 1 1 0 di-siRNA (mM)
B
*
Minutes
PBS
(no siRNA)
0 0 0 0 0 Ca2+ (mM)
4 8 12 16 32 Mg2+ (mM)
Divalent Cation Concentrations in aCSF Buffer
C
Minutes
PBS
(no siRNA)
16 15 14 13 12 11 10 Ca2+ (mM)
0 1 2 3 4 5 6 Mg2+ (mM)
Divalent Cation Concentrations in aCSF Buffer

**Figure S4.** The acute neurotoxicity and divalent cation content in aCSF buffer influence the time it takes mice to right themselves following oligonucleotide injections ICV. **(A)** Mice injected with 1mM (225μg/10μL total) di-siRNA$^{HTT}$ in aCSF with incrementally increased concentrations of $Ca^{2+}$ were sternal within similar times of the 1XPBS controls (ns). Excess $Ca^{2+}$ in aCSF without di-siRNA$^{HTT}$ delivered ICV resulted in significantly longer time taken for mice to be sternal compared to (**p=0.0021) and the balanced di-siRNA$^{HTT}$ aCSF injections (**p=0.0029 and *p=0.0349). Each data point represents one mouse (n=2-6); data were analyzed using one-way ANOVA followed by Tukey's multiple comparisons test**.** **(B)** Mice injected with 1mM (225μg/10μL total) di-siRNA$^{HTT}$ in aCSF with lower concentrations of $Mg^{2+}$ were sternal and experienced adverse events significantly quicker than 1XPBS controls (*p=0.0122 and *p=0.0349). There was no significant difference in the timing for mice injected with 12mM, 16mM, or 32mM $Mg^{2+}$. Each data point represents one mouse (n=4-6); data were analyzed using one-way ANOVA followed by Tukey's multiple comparisons test**.** **(C)** Mice injected with aCSF containing 16mM total divalent cations in a variety of ratios were sternal within the same time as 1XPBS controls. Each data point represents one mouse (n=4-6). No significance: data were analyzed using one-way ANOVA followed by Tukey's multiple comparisons test**.**

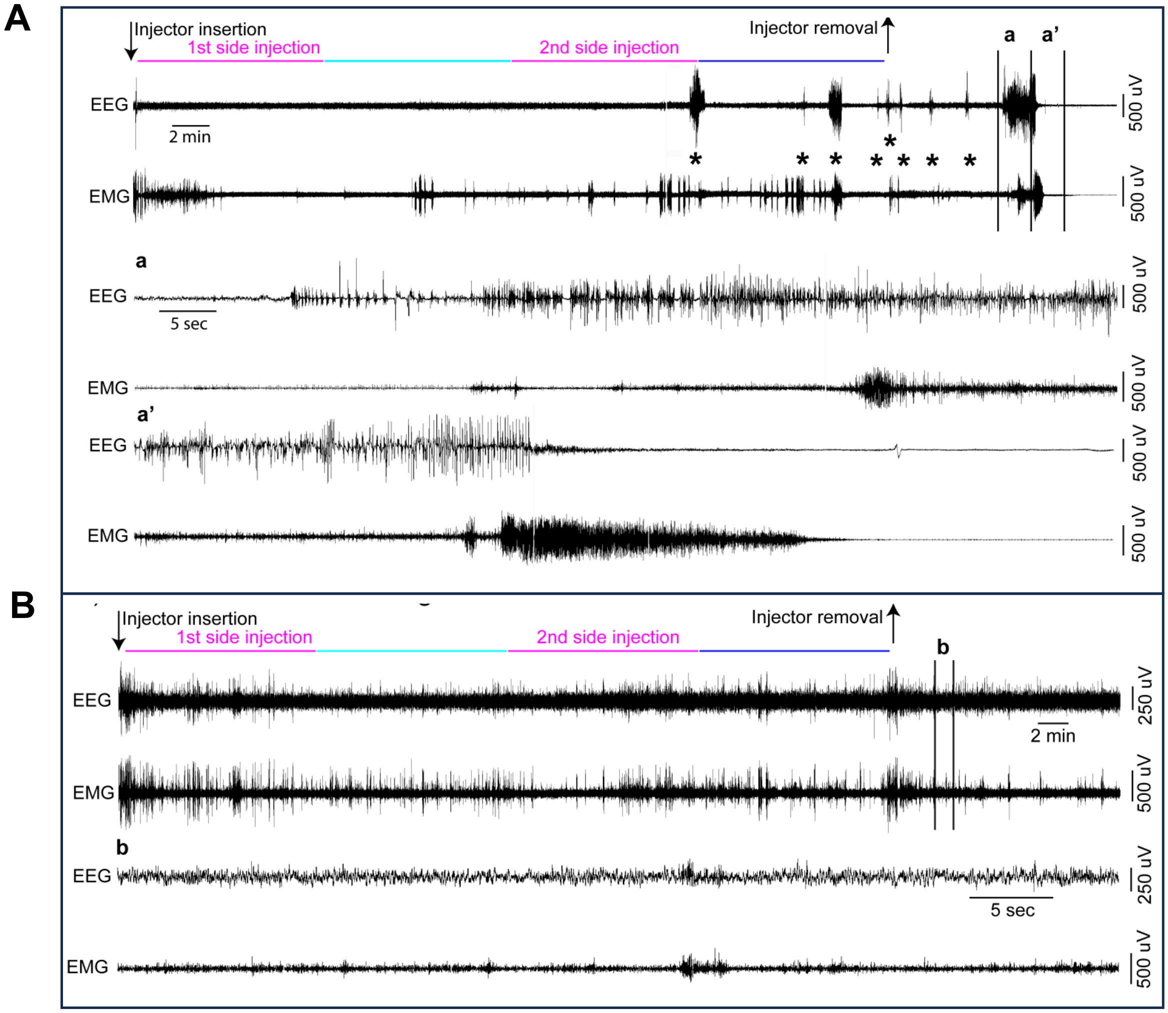


**Figure S5.** EEG/EMG examples from a freely moving mouse receiving bi-lateral ICV injection of 0mM (vehicle control) and 1mM (225μg) di-siRNA$^{HTT}$ in 10μL 1XPBS **(A)** or aCSF+ (**B**)**.** The top left arrow indicates the insertion of the injector in the mouse guide cannula. The compounds were first injected in one of the two lateral ventricles and then in the second lateral ventricle (5μL in 10 minutes, pink lines). Ten minutes separated the two side injections (light blue line). Following the end of the second side injection, the injector was left in place for an additional 10 minutes (dark blue line). The right arrow indicates the removal of the injector from the mouse guide cannula. (**a**, **a')** zoom into the portion of **(A)** top EEG/EMG delineated by the vertical bars. Note that the mouse injected with 1M (225μg) di-siRNA$^{HTT}$ in 10μL 1XPBS displays multiple seizures, characterized by high amplitude EEG waves and intense EMG activity, starting around the beginning of the second side injection. Interestingly, about 10 minutes following the injector removal, the mouse enters a prolonged seizure **(a-a')** that results in death, characterized by the absence of an EEG/EMG signal. Asterisks (*) denote seizures across the EEG example. (**b**) zoom into the portion of **(B)** top EEG-EMG delineated by the vertical bars. The mouse injected with 20nmol 1M (225μg) di-siRNA$^{HTT}$ in 10μL aCSF+ displays normal EEG-EMG activity during and after the ICV injection.

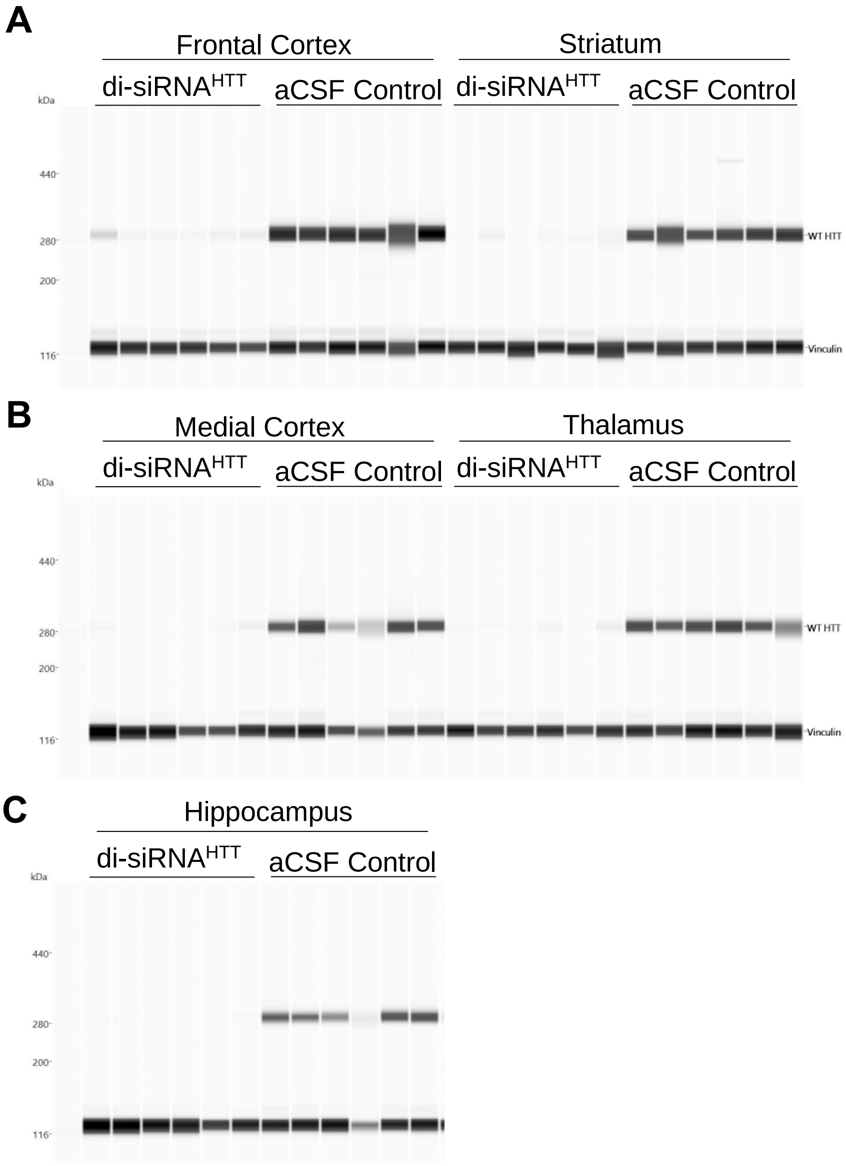


**Figure S6.** Raw western blot (ProteinSimple) of wild-type FVB/NJ mice treated with 1mM (225μg/10μL total) di-siRNA$^{HTT}$ in aCSF+ or aCSF control (Figure 5B). The level of wild-type mouse HTT protein was evaluated in the frontal cortex, striatum **(A)**, medial cortex, thalamus **(B)**, and hippocampus **(C)** two months following bilateral ICV injections (n=6).

A

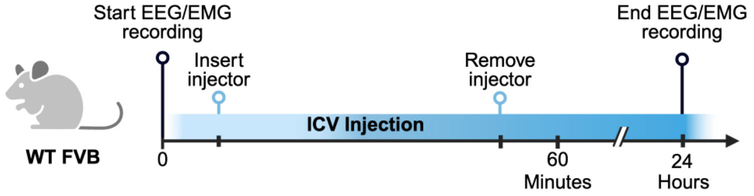


B

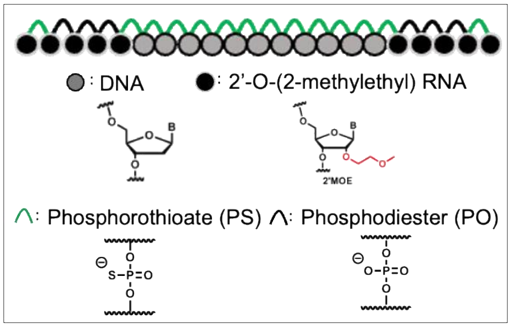


C

| Injection | Buffer | Seizure |
|---|---|---|
| 1mM ASO | 1XPBS | 0/6 |
| 1mM ASO | aCSF+ | 0/6 |

D

E

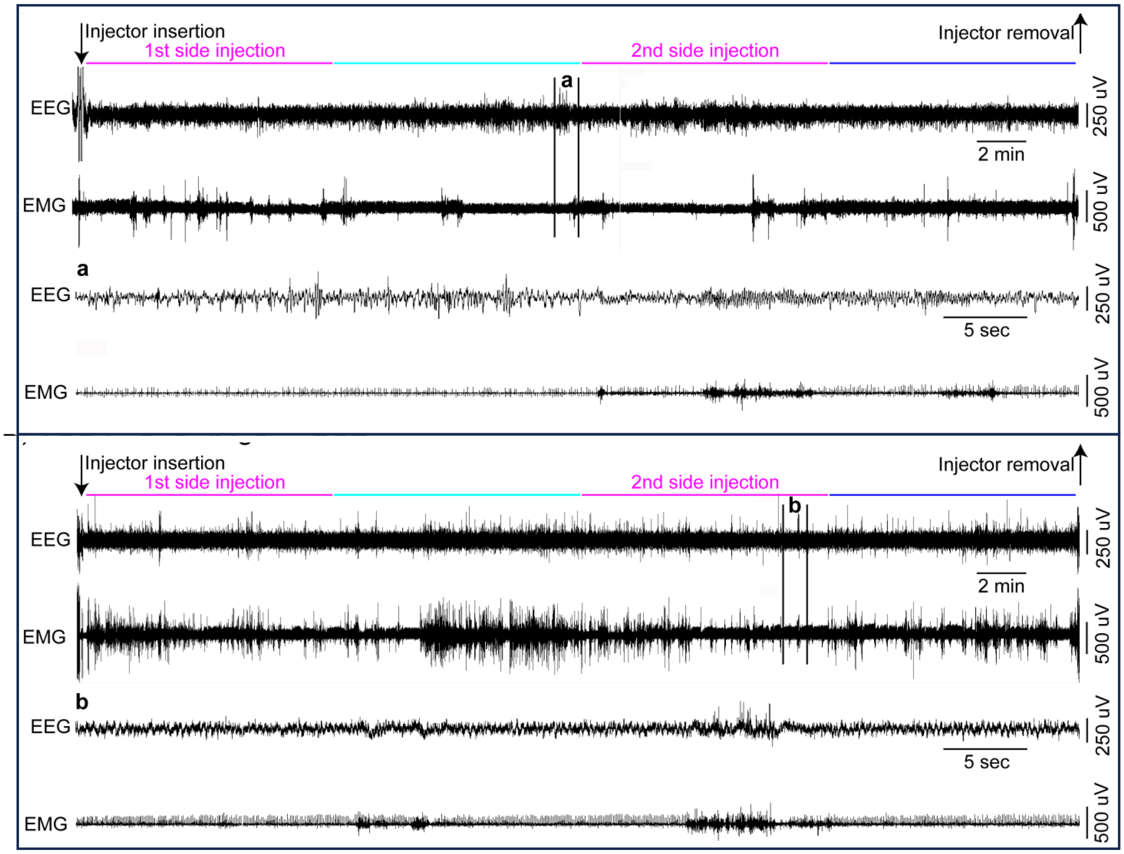

**Figure S7.** Electrophysiology confirmed the safety of a mixed PO/PS backbone ASO in awake mice. **(A)** Timeline showing ICV injection procedure and EEG/EMG recording for wild-type mice injected bilateral ICV with oligonucleotides awake. (Created with BioRender.com) **(B)** The chemical structure of the mixed backbone ASO (Tominersen[1]) used in this figure. **(C)** Table outlining the number of wild-type mice with seizure responses recorded by EEG/EMG following ICV injections of 3.2mM (225μg) mixed PO/PS ASO in 10μL 1XPBS or aCSF+ (n=6 per buffer condition). **(D-E)** EEG/EMG examples from a freely moving mouse receiving bi-lateral ICV injection of 225μg ASO in 10μL 1XPBS **(D)** or aCSF+ **(E).** The top left arrow indicates the insertion of the injector in the mouse guide cannula. The compounds were first injected in one of the two lateral ventricles and then in the second lateral ventricle (5μL in 10 minutes, pink lines). Ten minutes separated the two side injections (light blue line). Following the end of the second side injection, the injector was left in place for an additional 10 minutes (dark blue line). The right arrow indicates the removal of the injector from the mouse guide cannula. (**a**) Zoom into the portion of **(D)** EEG-EMG example, delineated by the vertical bars. (**b**) Zoom into the portion of **(E)** top EEG-EMG delineated by the vertical bars. Wild-type mice injected with 225μg ASO in 1XPBS and aCSF+ display normal EEG-EMG activity during the ICV injection **(a, b).**

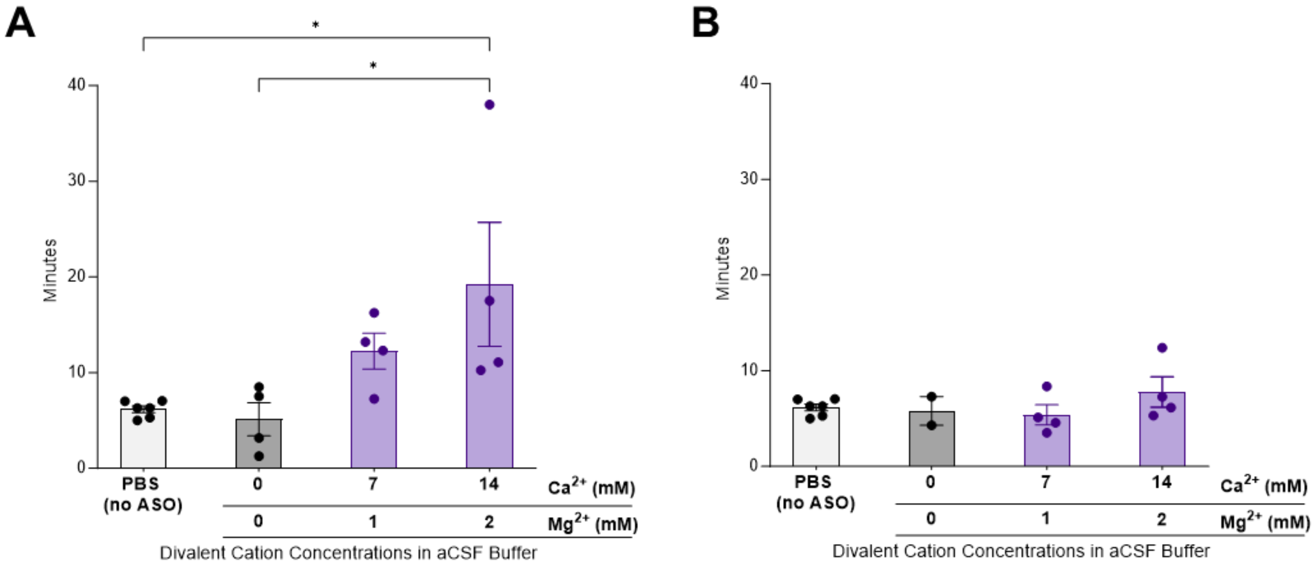


**Figure S8.** The acute neurotoxicity and divalent cation content in aCSF buffer influence the time it takes mice to right themselves following ICV injections of ASOs. **(A)** Mice injected with 225μg/10μL of full PS ASO in aCSF with 7:1 aCSF took approximately the same amount of time to be sternal compared to the full PS ASO delivered in 1XPBS and the 1X PBS vehicle control (ns). Mice injected with 225μg/10μL of full PS ASO in aCSF in 14:2 aCSF took a longer amount of time to be sternal compared to the full PS ASO delivered in 1XPBS and the 1X PBS vehicle control (*p=0.0329 and *p=0.0358). Each data point represents one mouse (n=4-6); data were analyzed using one-way ANOVA followed by Tukey's multiple comparisons test**. (B)** Mice injected with 225μg/10μL mixed PO/PS ASO in 1XPBS, 7:1 aCSF, and 14:2 aCSF were sternal within similar times of the mixed PO/PS delivered in 1XPBS and 1XPBS controls (ns). Each data point represents one mouse (n=2-6); data were analyzed using one-way ANOVA followed by Tukey's multiple comparisons test**.**

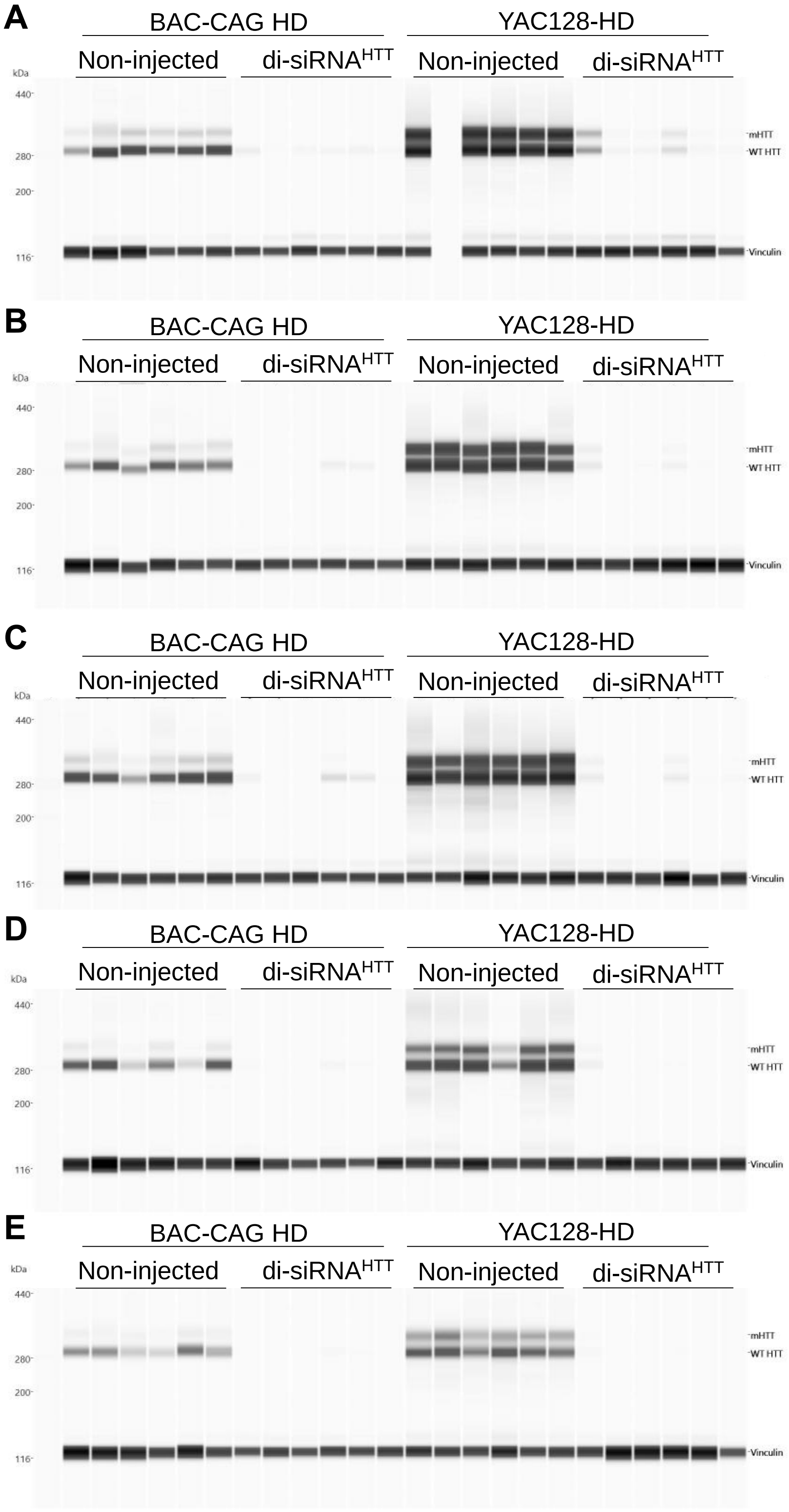

A
B
C
D
E
BAC-CAG HD
YAC128-HD
Non-injected
di-siRNAHTT
kDa
440
280
200
116
mHTT
WT HTT
Vinculin

**Figure S9.** Raw western blots (ProteinSimple) of YAC128-HD[2] and BAC-CAG-HD[3] mice treated with 1mM (225μg/10μL total) di-siRNA$^{HTT}$ in aCSF+ or aCSF control **(Figure 8G-J).** The levels of wild-type mouse HTT protein and mutant human HTT protein were evaluated in the frontal cortex **(A)**, striatum **(B)**, medial cortex **(C)**, thalamus **(D)**, and hippocampus **(E)** one - month following bilateral ICV injections (n=6).

**Supplementary Video 1:**
Comparison of the acute tolerability between WT mice injected ICV with either the 1xPBS control (left cage) or 1mM (225ug) di-siRNA in 1XPBS (right cage). The mouse in the left cage is experiencing a normal recovery whereas the mouse in the right cage is experiencing observable seizures.

**Supplementary Video 2:**
Comparison of the acute tolerability between WT mice injected ICV with either the 1xPBS control (left 1mM (225ug) di-siRNA in aCSF+ (left cage) or 1XPBS (right cage). The mouse in the left cage is experiencing a normal recovery, whereas the mouse in the right cage is experiencing observable seizures.
